# Multi-faceted deregulation of gene expression and protein synthesis with age

**DOI:** 10.1101/2020.01.19.911404

**Authors:** Aleksandra S. Anisimova, Mark B. Meerson, Maxim V. Gerashchenko, Ivan V. Kulakovskiy, Sergey E. Dmitriev, Vadim N. Gladyshev

## Abstract

Protein synthesis represents a major metabolic activity of the cell. However, how it is affected by aging and how this in turn impacts cell function remains largely unexplored. To address this question, herein we characterized age-related changes in both the transcriptome and translatome of mouse tissues over the entire lifespan. Expression of the majority of differentially expressed genes followed a U-shaped curve with the turning point around 3-months-old. We showed that transcriptome changes govern changes in the translatome and are associated with altered expression of genes involved in inflammation, extracellular matrix and lipid metabolism. We also identified genes that may serve as candidate biomarkers of aging. At the translational level, we uncovered sustained down-regulation of a set of 5’ terminal oligopyrimidine (5’TOP) transcripts encoding protein synthesis and ribosome biogenesis machinery and regulated by the mTOR pathway. For many of them, ribosome occupancy dropped 3-fold or even more. Moreover, with age, ribosome coverage gradually decreased in the vicinity of start codons and increased near stop codons, revealing complex age-related changes in the translation process. Taken together, our results reveal systematic and multi-dimensional deregulation in protein synthesis, showing how this major cellular process declines with age.

---

Aging is associated with a gradual decline of organismal function and fitness, which in turn is tightly linked with changes in the proteome. The balance between protein synthesis and degradation, proteostasis, and proper protein quality control are required to maintain cell homeostasis (1–3). Among other processes, molecular damage accumulating in cells with age influences the proteome, the endpoint of gene expression (4). Additionally, with advancing age, protein damage manifests itself in the form of post-translational modifications such as oxidation and glycation, impairing function; damaged proteins are also prone to form toxic oligomers and insoluble aggregates. In fact, disruption of proteostasis is a well-known cause of aging-associated diseases. However, this age-related dysfunction is also believed to influence organisms in a systemic and chronic way, decreasing their stress resistance and the ability to clear misfolded proteins (1). Indeed, compared to their closest relatives, long-lived species exhibit an increased proteome stability (5) and resistance to protein oxidation (6). Protein turnover and quality control include several distinct but tightly connected biological processes and can be conditionally divided into the phases of protein synthesis, folding, activity, post-translation regulation, and degradation.

## INTRODUCTION

Among them, only folding and degradation are relatively well understood in the context of aging, and their impairment indeed explains some aging-related pathologies (1). On the other hand, protein synthesis changes with age remain notably less explored (7, 8).

In the second half of the 20th century, several studies showed, by studying various species, that the overall rate of protein synthesis, activity, concentration of elongation and initiation factors, and tRNA aminoacylation levels decrease with age (9). Recent studies in mammals are also in agreement with the idea of age-related decline in overall translation rate. In particular, it was shown that the levels of total mRNA, as well as the expression of RNA polymerase I, eIF2Bε and eEF2, decrease with age in rat tissues (10). Increased promoter methylation in ribosomal RNA genes and decreased ribosomal RNA concentration during aging were also reported (11). In addition, down-regulation of translation with age was confirmed *in vivo* in the sheep (12) as well as in replicatively old yeast (13). Recently, analyses of liver and brain of 6- and 24-month old rats revealed age-related translatome changes in mammals (14).

The role of protein synthesis in aging is further supported by an indirect evidence. The decreased rate of protein synthesis generally leads to increased lifespan of animals (reviewed in (7)). For instance, knockout or knockdown of several translation machinery components in worms significantly increased average lifespan and accelerated the effects of life-extending mutations (15). In addition, overexpression of translation initiation repressor 4E-BP1 increased lifespan (16) and mediated lifespan extension effects of dietary restriction in fruit flies (17), whereas inhibition of biogenesis of the 60S ribosomal subunit prolonged lifespan of yeast (18). Notably, most of the interventions known to extend lifespan are associated with suppression of metabolism (e.g. caloric restriction) or inhibition of nutrient signaling (e.g. rapamycin), regulating protein synthesis and biosynthesis of translation machinery components (19). At least in part, this can be explained by the reduced load on the protein quality control machinery and decreased energy use (20). Moreover, increased fidelity of translation and decreased protein turnover rate were recently found to be associated with longevity; e.g. the naked mole-rat, an animal with extreme lifespan compared to its rodent relatives, possesses a highly accurate translation apparatus (21–23).

Despite a growing number of studies on protein synthesis alterations with age in mammals, previous research did not address translatome changes with the temporal and quantitative resolution sufficient to reveal principles of protein synthesis alteration with age at the whole-transcriptome scale. In this study, we took advantage of a combination of ribosome profiling (Ribo-Seq) (24) and RNA-Seq to characterize age-related changes in protein synthesis throughout the entire lifespan, focusing on liver and kidney of mice. This approach supported the identification of functional groups of genes exhibiting age-related changes in transcription and/or translation. Interestingly, dozens of transcripts encoding ribosome biogenesis and protein synthesis machinery components were specifically down-regulated with age at the translational level, consistent with the decline in protein synthesis with age. Ribo-Seq analyses also revealed a transcriptome-wide redistribution of ribosome coverage from the beginning to the end of mRNA coding regions as well as other features associated with complex and multi-factorial deregulation of protein synthesis with age.

## MATERIALS AND METHODS

### Tissue collection and lysis

Tissue samples were collected from male C57BL/6 mice of indicated ages from the NIA Aged Rodent Colony, as described in (25). Liver and kidney samples were sliced and frozen in liquid N_2_ and stored at −80°C. Tissue samples (~55 mg and ~75 mg for liver and kidney, respectively) were used for subsequent analyses. Tissues were lysed as described previously (4). After homogenization and centrifugation, 250 μl of lysate were supplied with 20 Units of SUPERase-In RNase inhibitor and taken for RNA-Seq library preparation, and the 500 μl of lysate was brought to 1 ml with lysis buffer and taken for Ribo-Seq library preparation.

### Ribosome profiling sequencing library preparation

Ribosome profiling libraries were prepared as described previously (4) with the modification in RNase digestion as indicated below. RNA digestion of lysates was performed for 1 hour with the mixture of 2000 Units of RNase T1 (Epicentre) and 300 Units of RNase S7 (Roche/Sigma). After 30 minutes of incubation, 0.8 mg heparin was added to inhibit all RNases except for RNase T1. After digestion, lysates were supplied with 80 Units of SUPERase-In RNase inhibitor.

### Transcriptome library preparation and sequencing

Total RNA was isolated from 250 μl of lysate with 750 μl of TRIzol LS Reagent and treated with RQ1 RNase-Free DNase (1 Unit for 1 μg of total RNA) for 30 minutes at 37°C with subsequent water saturated acidic phenol extraction and precipitation with ethanol (with the addition of 1/100 volume of glycogen RNA grade). 500 ng of DNase I treated total RNA was depleted of ribosomal RNA with NEBNext^®^ rRNA Depletion Kit (Human/Mouse/Rat) (#E6310) and used for transcriptome library preparation with NEBNext^®^ Ultra™ II Directional RNA Library Prep Kit for Illumina^®^ (#E7760). Both ribosome profiling and transcriptome libraries were sequenced on the Illumina NextSeq 500/550 system (Genome Sequencing Research and Education Center, Faculty of Bioengineering and Bioinformatics, Lomonosov Moscow State University).

### Ribo-Seq and RNA-Seq sequencing, data processing and bioinformatic analyses

The bioinformatic analysis of the Ribo-Seq and RNA-Seq sequencing data is described in detail in Supplemental methods. Briefly, the Ribo-Seq and RNA-Seq reads were aligned to the mouse transcriptome and genome assemblies (mm10, GRCm38.p5) using GENCODE M13 annotation. Transcriptome alignment was used to analyze ribosomal coverage changes and build metagene profiles. RNA-Seq and Ribo-Seq data were RLE-normalized, separately for kidney and liver, and transformed to CPM (counts per million). Differential expression was analyzed with the generalized linear model of the edgeR package. Both paired age comparison and linear model describing linear changes of gene coverage with age were used in the study.

## RESULTS

### Gene expression changes in mouse liver and kidney reflect age-related dysfunction

To characterize age-related changes in protein synthesis, we subjected mouse tissue samples to both transcriptome sequencing and ribosome profiling (Table S1), as this approach allows to separate the contribution of transcription and translation processes. Liver samples were collected from male mice representing six age groups (1, 3, 11, 20, 26 and 32 months old), and kidney samples from three age groups (3, 20 and 32 months old) (Fig. 1A). This broad range of ages was chosen to cover the entire adult mouse lifespan, from the final stages of development (1 month) to very advanced ages (32 months).

**Figure 1.**
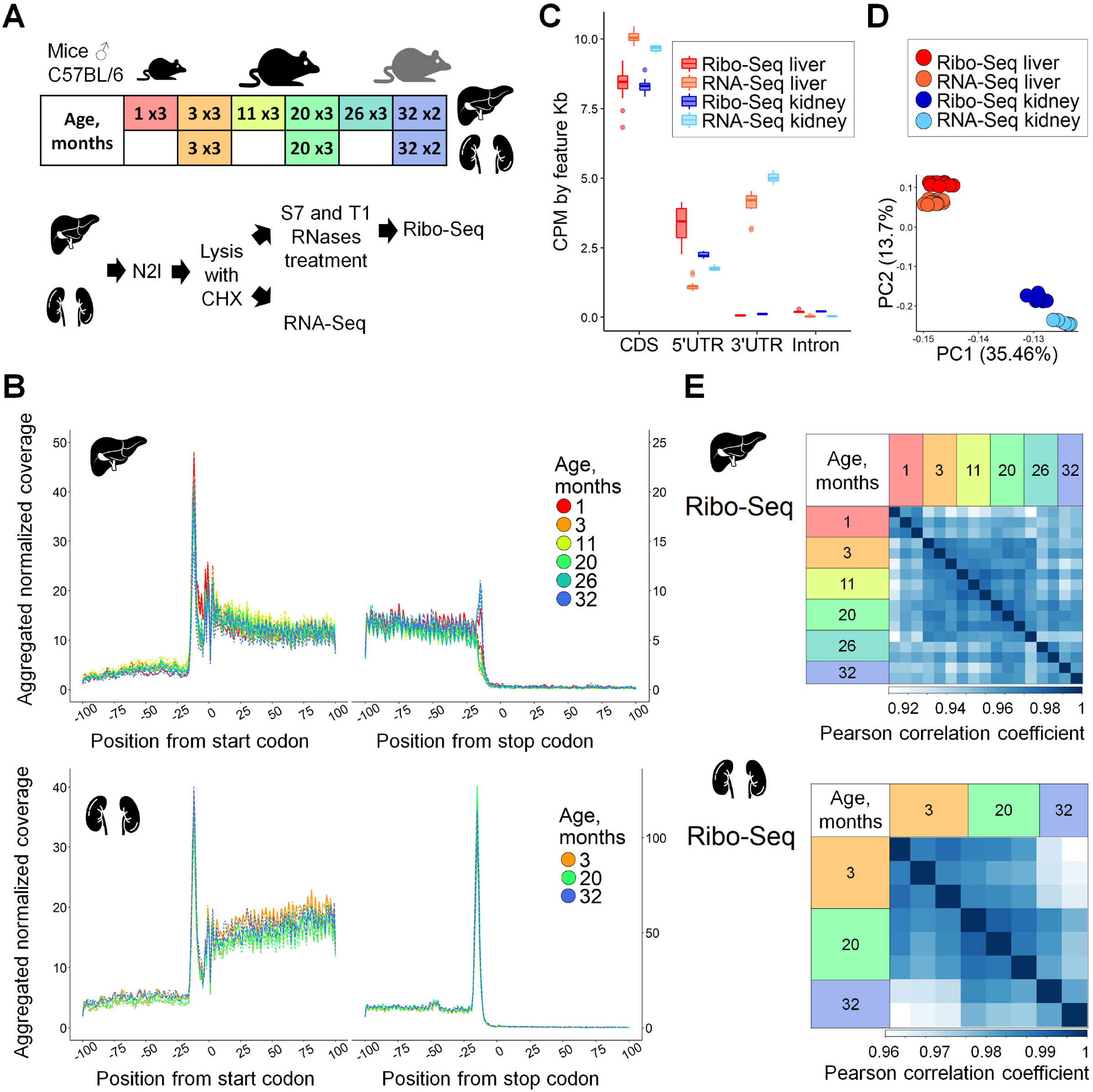
Ribo-Seq and RNA-Seq of aging mouse liver and kidney. (A) Overview of experimental design. Mouse livers representing six age groups (1-, 3-, 11-, 20-, 26- and 32-month-old) and kidneys representing three age groups (3-, 20- and 32-month-old) were used. For each age, three biological replicates were prepared (three C57BL/6 male mice), except for the 32-month group (two mice). Ribo-Seq and RNA-Seq libraries were prepared from the same cytoplasmic cell lysate. (B) Metagene profiles of ribosomal footprint 5’ends in 200 nt windows centered at start and stop codons built for 2,920 and 4,566 transcripts for liver and kidney, respectively. For each transcript, raw Ribo-Seq coverage was normalized to the sum of transcript coverage divided by its length. Normalized transcript coverage in the window was then aggregated for all selected transcripts. (C) Distribution of Ribo-Seq and RNA-Seq coverage in different gene regions. (D) Principal component analysis (PCA) of 8,562 genes in Ribo-Seq and RNA-Seq datasets of mouse liver and kidney. (E) Heatmaps of Pearson correlation coefficients for replicates of mouse liver and kidney analyzed by Ribo-Seq. For PCA and calculation of Pearson correlation coefficients, and further in the study, Ribo-Seq was analyzed together with the RNA-Seq dataset, but separately for organs. In total, the number of genes covered in each sample was 8,992 in liver and 11,461 in kidney.

In our Ribo-Seq protocol, we applied the combination of T1 and S7 RNases, because it most efficiently converts polysomes to monosomes in mouse tissues while retaining ribosome integrity (Fig. S1A) (26). The resulting footprints displayed clear triplet periodicity (Fig. 1B), the mean read length of 28 nt (Fig. S1B) and a higher CDS and 5’ UTR coverage compared to 3’ UTRs and introns (Fig. 1C). In both organs, there were pronounced peaks at start codons, while the peak at stop codons was more pronounced in kidney, in accordance with previous observations (27, 28). Principal component analysis (PCA) separated Ribo- and RNA-Seq samples by four groups according to their organ of origin and sequencing method (Fig. 1D). Expression profiles were highly reproducible across the replicates (Figs. 1E and S1C), especially within each age group, in both liver and kidney, reflecting the fact that gene expression profiles between individual mice within an age group are more similar than those observed across different ages. Samples from 1-month-old mice formed a separate group with high correlation across the replicates in both Ribo-Seq and RNA-Seq data (Fig. 1E and S1C), whereas for the older mice we detected a slight increase in gene expression variance with age (Fig. 1E and S1C), in agreement with previous reports (29).

We further assessed Ribo-Seq-based temporal patterns of gene expression in mouse liver (Figs. 2A and 2B) and kidney (Figs. 2C and 2D). Here, Ribo-Seq was not normalized to transcriptome coverage, therefore reflecting gene expression with respect to both transcription and translation changes. As liver from 1-month-old mice differed from that of other ages (Fig. 1E), we used 3-month-old mice as the reference. We examined all samples in comparison to the reference points of respective organs and selected genes that were differentially expressed in any of the comparisons (Fig. S3, Table S2). Most genes followed a U-shaped pattern of expression changes with the maximum or minimum at 3 months (Fig.2A). This turning point (30, 31) may reflect a transition from development to the adult state. Indeed, our analysis of genes with elevated expression in the 1-month old liver (compared to 3-month old) showed up-regulation of genes participating in urogenital and vascular system development, chromatin assembly, extracellular matrix organization, and amino-acid and nucleotide metabolism (Table S3).

**Figure 2.**
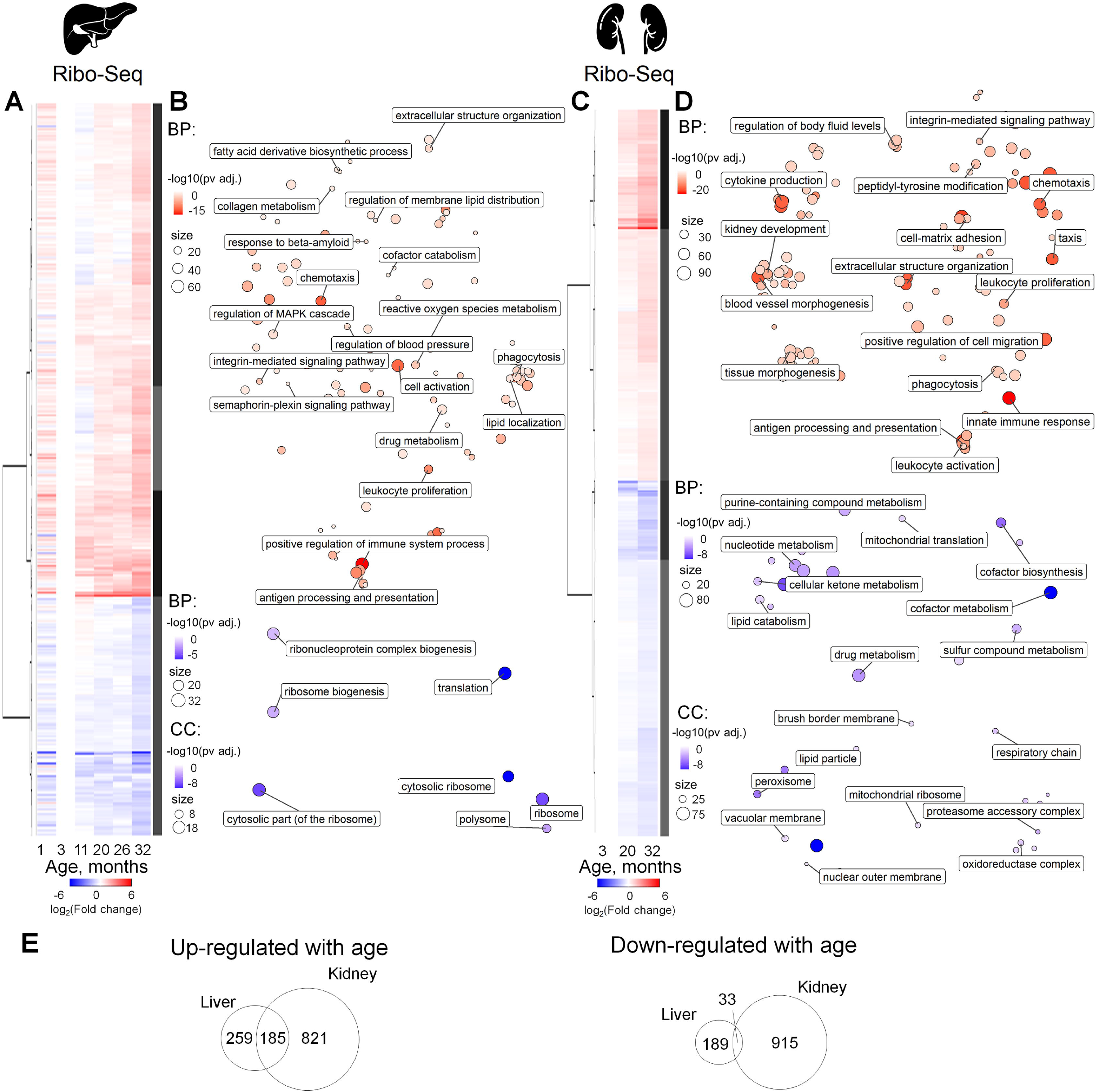
Expression profiling of aging mouse liver and kidney by Ribo-Seq. (A, C) Heatmap of age-related gene expression changes in liver (A) and kidney (C). For each age, differential expression, in comparison to that of 3-month-old mice, was calculated. For DE genes with adjusted P-value less than 0.05, at least in one age compared to 3 months old, log_2_(Fold change) values were clustered and presented on a heatmap (Table S2). The number of DE genes is summarized in Fig. S3. (B, D) GO BP (biological process) and GO CC (cellular compartment) functional enrichment of genes up-(red) or down-(blue) regulated with age in liver (B) and kidney (D) (Table S3). (E) Comparison of gene sets differentially expressed in liver and kidney. Venn diagrams show genes up- or down-regulated with age according to Ribo-Seq data.

Functional patterns of Ribo-Seq changes (Fig. 2E), as well as changes in the expression of particular genes (Fig. S2), were somewhat similar in liver and kidney, especially for genes up-regulated with age. We observed a robust increase in the expression of inflammation and immune system genes, reflecting “inflammaging” – chronic inflammation that progresses with age (reviewed in (30)). Common markers of senescent cells - lysosomal proteins (31), were up-regulated in both tissues. Five genes encoding inflammatory and lysosomal proteins (*Ctss, C1qa, C1qb, C1qc*, and *Laptm5*) were up-regulated in both liver and kidney in our study as well as several other aging transcriptome and translatome datasets in different organs and species (14, 32, 33), suggesting that they may be considered to be reliable biomarkers of aging.

In contrast to inflammation, mitochondrial function is known to decline with age (reviewed in (34)). Indeed, in kidney we observed a gradually decreased expression of nuclear genes coding for mitochondrial proteins, although it was less pronounced in liver (yet many genes followed the pattern, e.g. *Uqcc2, Fxn, Mrps16, Mrpl9, Mrpl30, Mrpl54, Mterf4, Atp5k, Aadat, Mtch2*). Expression of genes participating in redox homeostasis was changed in both liver and kidney, consistent with the role of oxidative stress in aging (reviewed in (35)). Some of these genes were down-regulated in kidney and liver (*Pex16, Sod1*), whereas other genes were up-regulated (*Gpx3, Gsta2* - in both liver and kidney; *Gstt3, Sod3* – in liver; *Gstt1* – in kidney). Other functional groups of genes with increased expression in both organs were related to regulation of blood pressure and precursors of amyloid proteins (*Prnp* - in liver and kidney; *App, Aplp2* – in liver).

Genes downregulated in kidney included those participating in the response to glucocorticoid hormones, cofactor biosynthesis, and lipid metabolism. Notably, 21 genes encoding peroxisomal components showed decreased expression with age in kidney, in agreement with association of age-related alterations in lipid metabolism with renal disorders (36).

The majority of age-related changes in the translatome were correlated with those in the transcriptome in both organs, suggesting that the primary changes of gene expression with age occur at the transcriptional level (Fig. S3). Therefore, we searched for putative transcription factors (TFs) regulating genes up- or down-regulated with increased age (Table 1, Table S4). This analysis revealed RelA (an NF-κB p65 subunit) and Spi-B (lymphocyte-specific transcription activator) which may activate transcription of their targets with aging in both liver and kidney. Most of their targets up-regulated with age participate in inflammatory response and immune processes. In addition, 55 out of 217 RelA targets up-regulated in kidney were found to be shared with another transcription factor, Jun (an AP-1 transcription factor subunit), which in cooperation with NF-κB promotes cell survival (37). In turn, shared targets of Jun and Smad3 were also enriched for genes with an age-related increase in expression, reflecting the response to pro-apoptotic cytokine TGFβ (38). Additional factors that can participate in transcriptional activation of immunity-related genes with age were MafB in liver and Irf3 and Stat5a in kidney. Targets of peroxisome proliferator-activated receptor α, PPARα (alone or together with retinoic acid receptor RXRα) were enriched for up-regulated lipid metabolism genes in kidney. The PPARα/RXRα dimer activates transcription of genes participating in fatty acid oxidation and catabolism (39). Moreover, activation of PPARα represses NF-κB function, and the reduction in PPARα levels is associated with aging (40). *Ppara* itself is among the Foxa3 transcription factor targets, which was enriched in kidney genes up-regulated with age.

**Table 1.** Transcription factors with binding sites enriched in promoters of genes up- or down-regulated with age.

| Gene Symbol | P-value adj* | Odds ratio | Number of targets |
| --- | --- | --- | --- |
| Potential regulators up-regulated with age in liver |  |  |  |
| *Spib* | 9.17E-03 | 3.29 | 19 |
| *Rela* | 1.25E-02 | 1.57 | 109 |
| *Mafb* | 4.96E-02 | 1.84 | 42 |
| Potential regulators up-regulated with age in kidney |  |  |  |
| *Irf3* | 1.87E-06 | 2.43 | 66 |
| *Msx1* | 4.67E-04 | 1.43 | 261 |
| *Rela* | 7.64E-04 | 1.44 | 217 |
| *Spib* | 1.97E-02 | 2.29 | 28 |
| *Ppara* | 2.34E-02 | 1.49 | 101 |
| *Cebpa* | 4.41E-02 | 1.27 | 254 |
| *Foxa3* | 4.41E-02 | 1.86 | 36 |
| *Jarid2* | 4.41E-02 | 1.23 | 535 |
| *Stat5a* | 4.64E-02 | 1.30 | 194 |
| Potential regulators down-regulated with age in kidney |  |  |  |
| *Hnf4a* | 1.55E-02 | 1.45 | 156 |
| *Arid1a* | 1.55E-02 | 1.35 | 244 |
| Interactions of two transcription factors |  |  |  |
| Potential regulators up-regulated with age in kidney |  |  |  |
| *Rela + Jun* | 1.10E-02 | 1.82 | 55 |
| *Ppara + Rxra* | 3.08E-02 | 1.56 | 77 |
| *Ppara + Gata4* | 3.08E-02 | 2.93 | 16 |
| *Msx1 + Tbp* | 3.20E-02 | 1.92 | 35 |
| *Jun + Smad3* | 3.20E-02 | 1.71 | 49 |
| *Stat5a + Esr1* | 3.20E-02 | 1.50 | 83 |

\* P-values of right-sided Fisher's exact test were adjusted with the Benjamini-Hochberg correction.

For down-regulated genes, we found two potential transcription factors involved in kidney, Arid1a and Hnf4a, regulating expression of genes responsible for lipid and cofactor metabolism

### Aging affects ribosome occupancy of specific groups of transcripts

Intriguingly, many down-regulated genes were related to protein synthesis, non-coding RNA metabolism, and ribosome biogenesis, at least in liver. Among them were the genes encoding numerous ribosomal proteins, translation factors, proteins involved in large (*Nol9, Nsa2*) and small (*Utp14a, Tsr1*) ribosome subunits biogenesis, RNA polymerase I components (*Cd3eap*), nuclear import and export proteins (*Nmd3, Ipo4, Sdad1*), RNA helicases (*Ddx1, Ddx17, Ddx21*), non-coding RNA processing nucleases (*Elac2*), aminoacyl-tRNA synthetases (*Mars, Qars, Farsb*), components of rRNA pseudouridylation complex (e.g. *Nhp2*) and RNA binding proteins with various functions (*Cirbp, Aimp1, Pa2g4, Rtraf*). In kidney, various genes related to protein synthesis also showed a decreased expression (e.g. *Eif3h, Eif4g3, Eif5, Eif5a, Dars, Sars, Ddx3x, Cirbp, Rtraf*). This functional group was not observed in previous age-related transcriptome studies, suggesting down-regulation specifically at the post-transcriptional level. To investigate this further, we analyzed contributions of transcription and translation by comparing age-related changes detected in RNA-Seq and Ribo-Seq data (Figs. 3 and S3).

**Figure 3.**
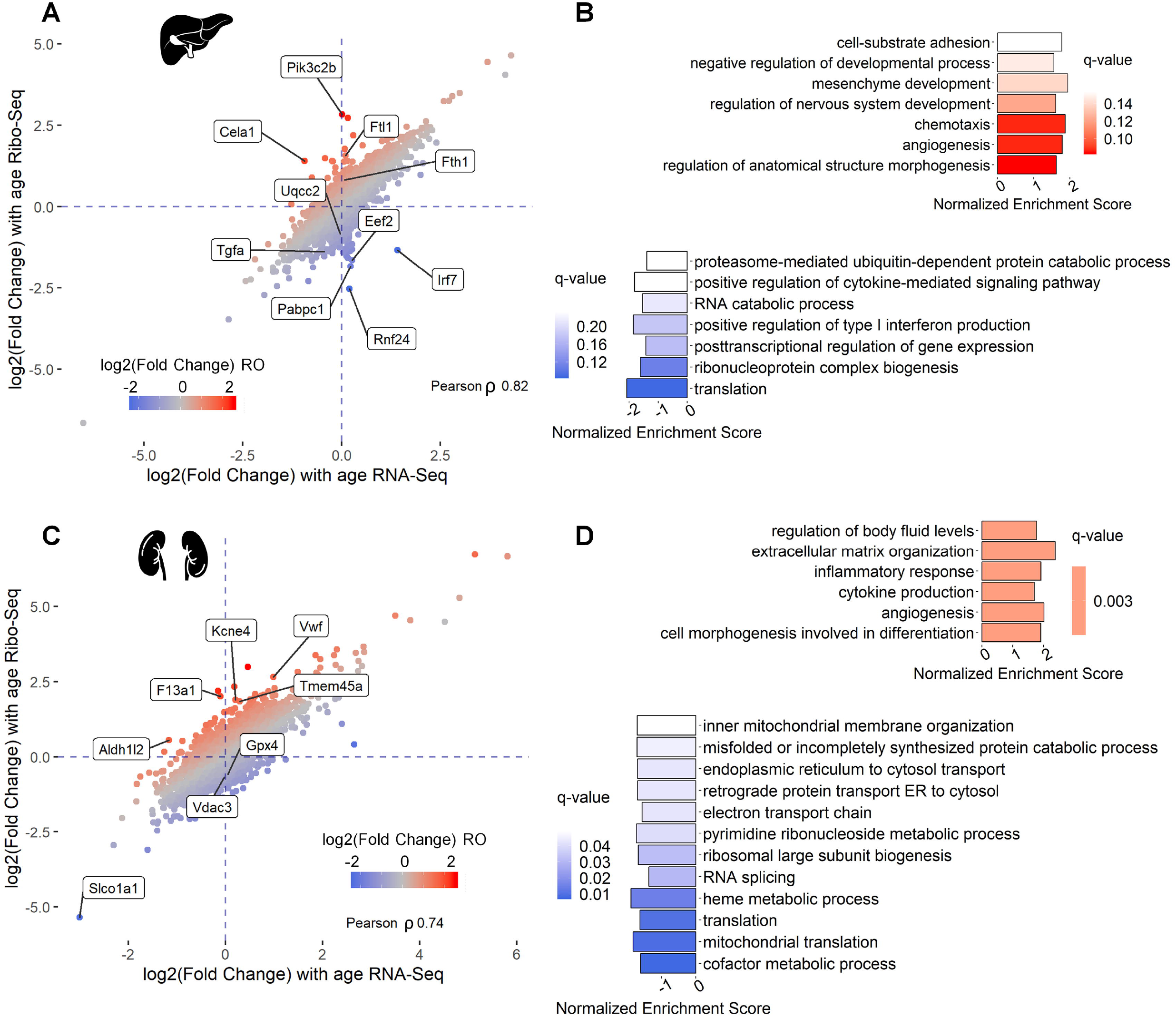
Age-dependent changes in ribosome occupancy of functional gene groups. (A, C) Comparison of transcriptional (RNA-Seq) and translational output (Ribo-Seq) for liver (A) and kidney (C) of 32-months-old mice vs. 3-months-old-mice (Table S2). (B, D) Functional groups of transcripts with age-related changes in RO presented as the results GSEA for genes sorted according to their RO changes, in liver (B) and kidney (D). Non-redundant GO BP terms with q-value less than 0.25 are shown (Table S3).

Together, Ribo-Seq and RNA-Seq allow decomposition of the contribution of transcription and translation to gene expression changes by analyzing ribosome occupancy (RO, number of ribosome footprints normalized to transcript abundance) of particular transcripts. RO is a proxy of translation efficiency (TE), since in most cases the more ribosomes translate the mRNA, the more products are made. Thus, we first compared age-dependent changes in gene expression identified by Ribo- and RNA-Seq in each organ (Figs. 3A and 3C). Even though age-related changes in gene expression were mostly driven by transcriptome changes, there were also distinct outliers, and almost all of them were differentially expressed at the translational and not the transcriptional level.

We performed gene set enrichment analysis (GSEA) of genes sorted by age-dependent RO fold changes (Figs. 3B and 3D, Table S3). In both organs, up-regulated genes represented the GO terms related to inflammation, development and differentiation of different cell types. The GO terms encompassing down-regulated genes in kidney included mitochondrial genes (components of the electron transport chain, mitochondrial translation, and membrane). Thus, several functional groups of genes indeed exhibited translational regulation affected by aging although there were no individual genes passing FDR adjusted P-value.

However, by examining the top and bottom of the ranked gene list, we found genes encoding functionally related products. Ferritin subunits Fth1 and Ftl1, known to be translationally regulated by an iron-responsive element in their 5’ UTRs (41), were both up-regulated in liver. Among the genes with the RO decreased with age, a striking example was interferon regulatory transcription factor 7 (Irf7), known to be translationally repressed through 4E-BPs (42). Its RO was ~6 times lower in old compared to young samples. The most prominent GO terms enriched in down-regulated genes in liver were related to protein biosynthesis: translation, ribosome biogenesis and post-transcriptional regulation (Figs. 3A and 3C).

The “translation” GO term included genes that were also down-regulated in kidney, although with lower significance. It could be that the observed changes in RO are linked to changes in transcript isoform abundance. To clarify this, we analyzed changes in transcript isoform abundance with age (Table S5) and compared them with RO changes of the corresponding genes. However, we detected neither significant switches of major transcript isoforms with age nor association between isoform abundance and RO changes (Figs. S4, S5).

### Translation of 5’ TOP transcripts encoding components of protein synthesis machinery is repressed with age

To assess possible mechanisms of the age-dependent translational decrease of transcripts encoding components of the protein biosynthesis machinery, we focused on top down-regulated transcripts from the GO term “translation” (GO:0006412). In liver, most of these genes had a uniform pattern of change: Ribo-Seq counts increased from 1 to 3 months and decreased gradually from 3 months to the oldest age, whereas RNA-Seq counts did not change with age or had a high level of variance (Fig. 4A). For example, genes coding for ribosomal proteins Rps5, Rps11, Rps25, Rps21, elongation factor Eef2 and Pabpc1 were among the top with the RO decreased in liver (Fig. 4B). Indeed, translation of most mTOR-sensitive transcripts and almost all 5’ TOP transcripts was down-regulated with age. Thus, the data suggest that the observed decline in the translation of transcripts encoding components of the protein synthesis apparatus could be explained by down-regulation of mTOR with age. Interestingly, we found that the abundance of mTOR mRNA itself was negatively associated with age according to our Ribo-Seq and RNA-Seq data (Fig. 4E), suggesting that during aging mTOR kinase can be regulated at the transcriptional level.

**Figure 4.**
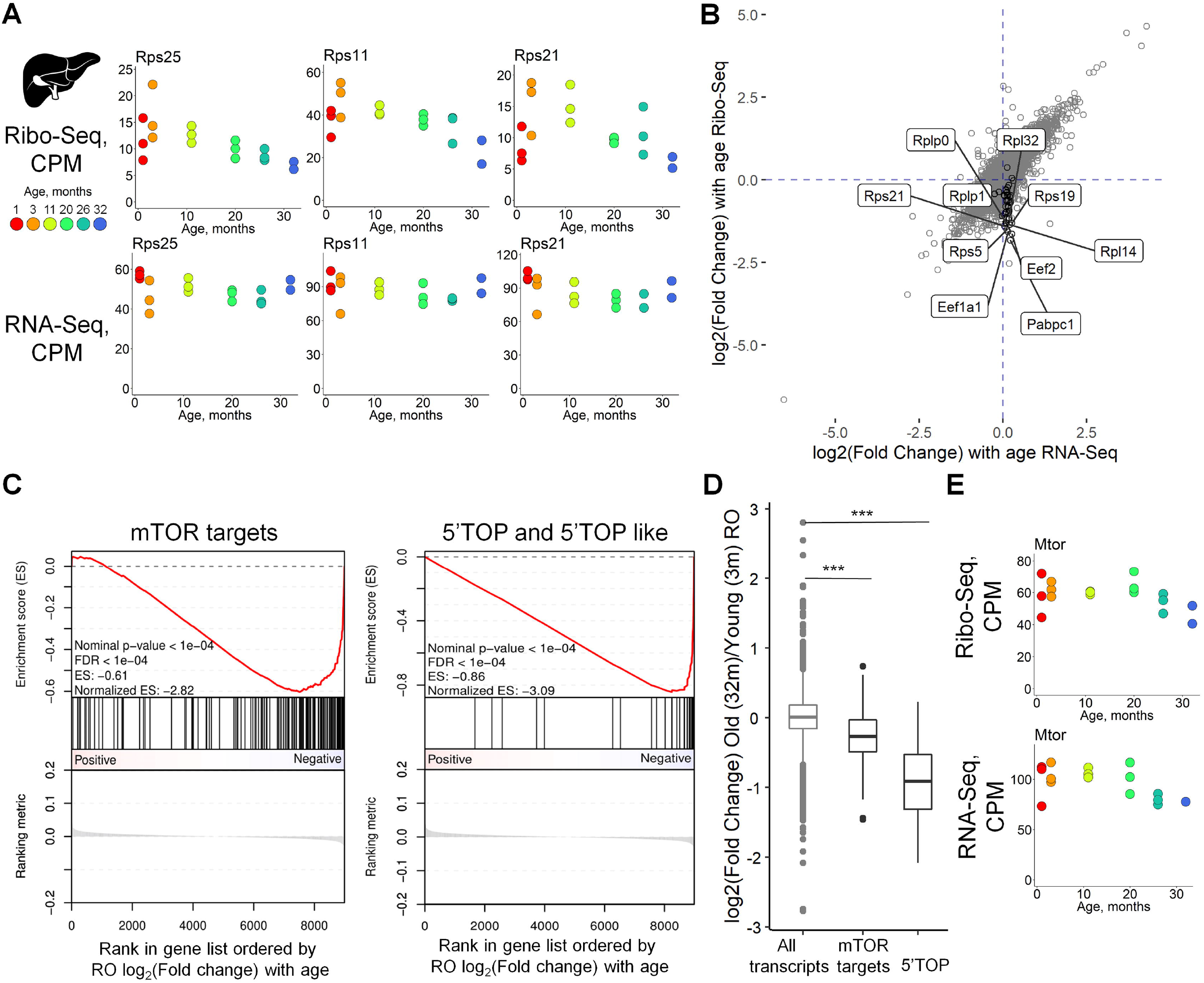
Decreased ribosome occupancy of transcripts encoding ribosomal and other translation-related proteins with age. (A) Examples of age-dependent changes in Ribo- and RNA-Seq counts of genes coding for proteins with functions associated with translation. (B) Comparison of transcriptome (RNA-Seq) and translation output (Ribo-Seq) of 32-months-old mice to 3-months-old-mice. Black dots represent 5’TOP genes. (C) GSEA of ribosome occupancy age-related changes (log2(Fold change), linear from 3- to 32 months old mice) in liver of 41 5’ TOP and 160 mTOR-sensitive genes (43). (D) Box plot showing distribution of mTOR regulated and 5’TOP genes ROs. Statistical significance was calculated with Mann-Whitney test. (E) Ribo- and RNA-Seq counts of Mtor gene.

In kidney, a similar but weaker pattern of changes was observed for transcripts coding for translation-related components (Fig. S6A). Although Ribo-Seq counts slightly decreased from 3 to 32 months, RNA-Seq did not change, or even increased, for most of these genes, thus compensating for the observed effect. This is probably why the translation GO term was not detected in the overall differential expression profile in kidney (Fig. 2A). Nevertheless, even with a similar pattern of change in translation-related transcripts and RO, the repertoire of these transcripts was not identical for liver and kidney (e.g. compare Figs. 4B and S6B, see also Table S6). We also observed down-regulation of 5’ TOP transcript translation with age (Fig. S6C) and a decrease in mTOR transcript levels (Fig. S6E), while in this case we did not find a pronounced decrease in RO of mTOR-sensitive transcripts (Fig. S6C).

### Redistribution of ribosome coverage towards the 3’ end of coding sequences with age

We constructed metagene profiles of ribosomal coverage in 100 nucleotide windows surrounding the start and stop codons of reliably expressed transcripts in liver and kidney (Fig. 5A). Strikingly, we observed an age-dependent decrease in ribosomal footprint density at the 5’ end of coding regions and an increase at its 3’ proximal part, in both organs. A change in ribosome distribution of a limited subset of mRNAs may strongly contribute to the observed pattern, but it may also be explained by minor changes in many mRNAs. To distinguish between these possibilities, we analyzed the positional profiles of ribosome footprints along individual transcripts, relative to the 3-month reference point. We split each transcript into a fixed set of segments (see Methods) and fitted a linear model of footprint coverage using a relative transcript coordinate as the predictor variable (Fig. 5B, Fig. S7; Supplementary Materials, Section 4). For each age, we found a significant shift of the distribution of linear regression slopes across transcript (compared to the younger age data using the sign test (Fig. S7)), whereas only a small number of transcripts showed ribosome coverage increases in the 5’ proximal part. Thus, the changes in ribosomal coverage along transcript reflect a global tendency of translation deregulation with aging. These age-dependent changes are illustrated in Fig. 5C by showing representative transcripts. The effect was most pronounced in the vicinity of start and stop codons, but also was detectable along whole transcripts (Fig. S7A). Overall, this redistribution pattern uncovers yet another layer of age-related translation deregulation.

**Figure 5.**
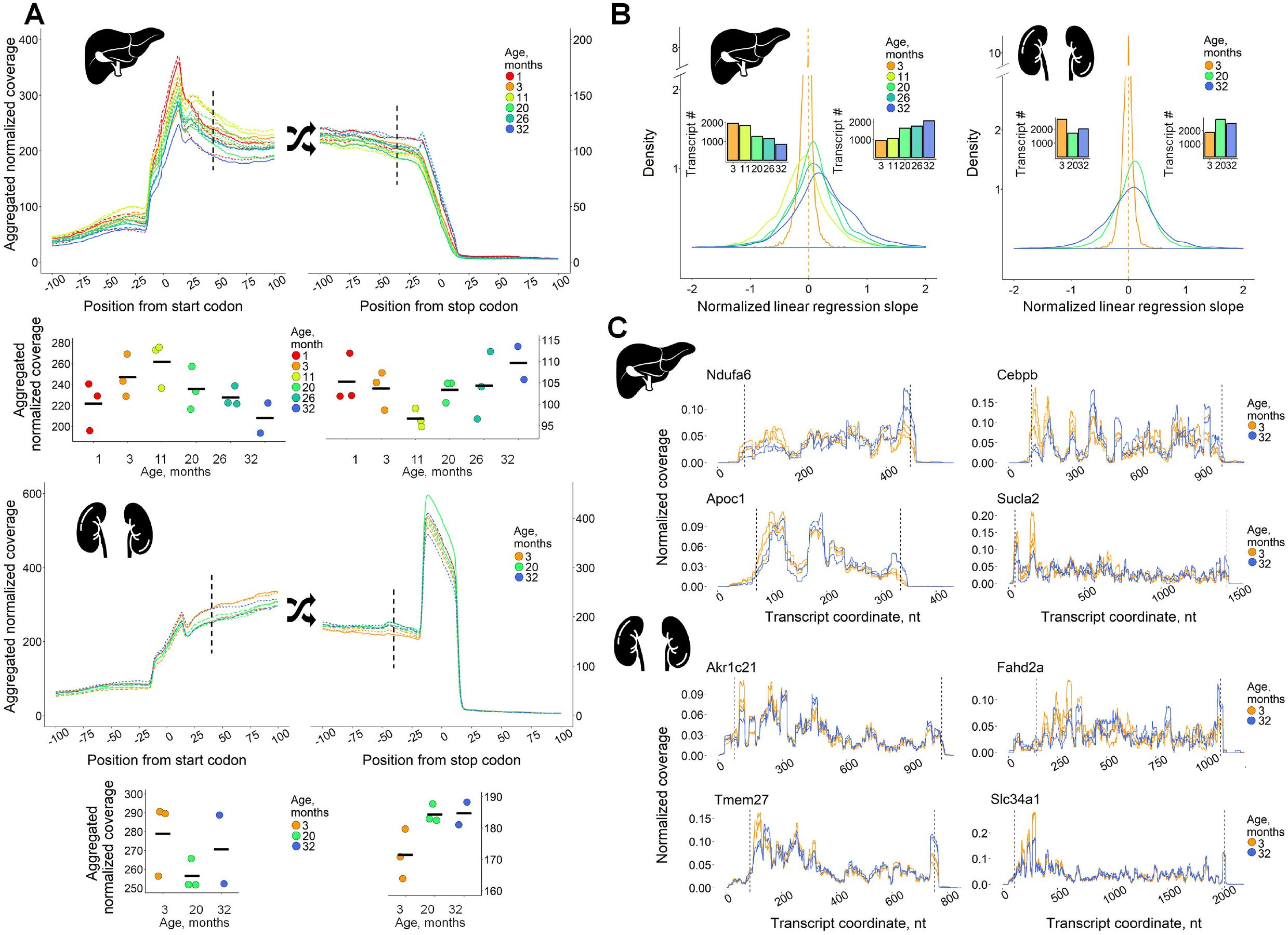
Gradual age-related rearrangement of ribosome footprints towards the 3’ end of coding sequence. (A) Metagene profiles of ribosomal coverage in the vicinity of start and stop codons (200 nt windows) of 2,920 and 4,566 transcripts for liver and kidney, respectively. Metagene coverage values at +42 nt position from start-codons and at - 42nt position from stop-codons are presented on separate graphs below the main graphs (coverage values in replicates, mean shown as horizontal lines). (B) Distribution of linear regression slopes for ribosome footprint profiles normalized to mean coverage at 3 months and smoothed with the relative transcript coordinate as the predictor variable (see Supplementary Materials, Section 4). Bar plots depict the number of transcripts with negative (left) and positive (right) slopes. (C) Representative transcripts exhibiting increased ribosome footprint coverage in liver and kidney with age. Dashed lines denote start and stop codons.

## DISCUSSION

Alteration of protein synthesis with age has been a contentious issue for some time (reviewed in (7)). Although the overall protein synthesis in mammals is thought to decrease with age, mechanistic details remained elusive. In this work, we applied Ribo-Seq and RNA-Seq to analyze age-related changes in the translatomes of mouse liver and kidney over the entire lifespan. This allowed us to characterize functional groups of genes, whose expression is altered with age at either transcription or translation levels, and to uncover candidate genes that may serve as markers of aging in examined tissues. In addition, we identified specific translational deregulation of 5’ TOP transcripts encoding components of protein synthesis machinery during aging. We also discovered that the positional profile of ribosomes along the transcript differs between young and old animals, reflecting a further layer of age-related changes in translation as animals age.

The gene expression changes that we observed (Fig. 2) were generally consistent with the previously described transcriptomic patterns during aging (e.g. (32, 33)), pointing to the major processes altered with age at the level of gene expression, such as inflammation, regulation of blood pressure, lipid and glucocorticoid biosynthesis, proteasomal protein degradation, mitochondrial activity and oxidative stress. A strong correlation between Ribo-Seq and RNA-Seq profiles as well as concordance with data reported in previous studies indicate that age-related changes in gene expression are manifested predominantly at the transcriptional level, at least in liver and kidney (Fig. 3).

Expression of the majority of genes differentially expressed with age followed a U-shaped curve with the turning point around 3 months of age. In previous studies, such a pattern was not observed, probably because age-related gene expression changes were examined either only at two time points (14, 32) or they did not include young animals (1 month old) (14, 32, 33). However, the U-shaped, “reversal” pattern of gene expression was previously reported for human (44, 45) and rat (46) brain, with the turning points at ~3.5 and ~20 years for humans and between 6-12 months for rats.

Apart from age-related changes in gene expression at the transcriptome level, we found a number of transcripts affected specifically at the level of translation. These include transcripts coding for mitochondrial proteins, immunity-related transcription factor Irf7, clinically-relevant hemostasis factors F13a1 and Vwf, ferroptosis-related peroxidase Gpx4 and two ferritin subunits, Fth1 and Ftl1. Additionally, genes from the GO terms related to development and cell differentiation were enriched among the genes with RO up-regulated with age in both organs. On other hand, we did not find association between isoform abundance and RO changes (Table S5, Figs. S4, S5), suggesting that alternative splicing is unlikely to be the source of the observed RO changes.

Most importantly, our analysis revealed a particular subset of transcripts, the 5’ TOP mRNAs encoding multiple components of protein synthesis machinery, whose translation efficiency is gradually decreased with age (Fig. 4). This group of transcripts is specifically regulated by the mTOR/4E-BP axis (47), a signaling pathway known to be associated with aging, lifespan control and longevity (for review, see (7)). Interestingly, we also observed down-regulation of mTOR mRNA abundance with age, suggesting the existence of a transcriptional component of mTOR regulation during aging (Fig. 4E). Although ribosome profiling provides expression data for protein-coding genes only, it is likely that rRNA and tRNA synthesis are also compromised during aging, as this expression is controlled by the mTOR pathway as well (48).

Thus, we showed that translation of mRNAs encoding protein synthesis machinery components is decreased with age in both liver and kidney. This pattern well correlates with the previously observed decline in overall protein synthesis with age (for review, see (7)). However, in contrast to rapidly proliferating cancer cells, where the mTOR-dependent transcripts constitute a major fraction of polysome-associated mRNAs, in our data, collected from terminally differentiated cells of mouse organs, 5’ TOP transcripts were not enriched among the highly translated transcripts. Therefore, their decreased association with polysomes is not supposed to significantly contribute to the overall decline in protein synthesis in old animals, but rather affects indirectly by altering translation machinery abundance and composition. Of note, the observed decrease in the overall expression of translation machinery components is most notable in liver, whereas in kidney changes in the translation rate of these transcripts are substantially compensated for at the transcriptional level (Fig. S6). We also cannot exclude a contribution of age-related changes in tissue composition, as whole tissue lysates were used for experimental analyses.

The described translational downregulation of transcripts encoding components of protein synthesis machinery was not detected in the pairwise comparison of gene expression in liver and brain of young and old rats, performed by an earlier ribosome profiling study (14). However, the findings are consistent with the translatome changes during yeast replicative aging (13). Interestingly, as the yeast transcripts encoding translation-related components do not exhibit 5’ TOP motifs (47) and thus are unlikely to be regulated by mTOR in the manner similar to the mammalian transcripts, in this case the reduction of both overall translation and ribosome protein synthesis is achieved by distinct mechanisms, i.e. activation of the GCN2/eIF2α regulatory pathway and elevated mRNA recruitment to P-bodies in aging cells (13). Interestingly, the GCN2/eIF2 axis was recently shown to regulate translation of 5’ TOP mRNAs also in mammals (48), so its possible link with the observed phenomenon is worthy of further investigation.

Another important observation in our study concerns changes in the metagene profile of ribosome coverage along coding regions (Fig. 5), pointing to a systemic alteration of translation with age. We suggest that these changes reflect the decrease in overall translation initiation efficiency caused by the observed down-regulation of the mTOR pathway (Fig. 4), as well as a possible decline in translation termination or ribosome recycling rate. Our model is based on the idea of the differential elongation speed in different parts of the coding sequence(49, 50). It is known that slow codons are distributed within transcripts in a non-random fashion and are particularly enriched in the region following the initiation site, while the distal parts of the coding regions have a much lower content of slow codons (49, 50). As a result, the first ~30-50 codons are usually translated more slowly, whereas the last ~50 codons are the fastest (49). Thus, under conditions when translation initiation is inhibited, ribosome density along the coding sequence should be redistributed from the 5’ proximal to the distal part of the transcript, while inhibition of termination and/or of ribosome recycling should increase the density in its 3’ proximal part. It should be noted, however, that elevated elongation speed and perhaps other factors could also lead to a similar pattern of metagene profile changes with age (51).

In summary, our results revealed previously unknown modes of translation deregulation with age. The decrease in translation rate may reflect an attempt of the cell to cope with the accumulation of damaged proteins or compensate for the deficit of energy with age (7). For many model organisms from yeast to primates, pharmacological, dietary, and genetic interventions reducing protein synthesis rate and inhibiting mTOR signaling have been shown to significantly increase lifespan (7, 8, 52). However, our study clearly shows that younger tissues are actually characterized by more active protein synthesis and elevated translation of mTOR-dependent mRNAs. Thus, returning the cell to a younger state should include renewal and re-activation of protein synthesis machinery, accompanied by simultaneous reinforcement of the cellular proteostasis network. As revealed by our study, age-related translation deregulation has many faces, together contributing to dysfunction in this most important cellular process during aging.

## DATA AVAILABILITY

All raw and processed sequencing data generated in this study have been submitted to the NCBI Gene Expression Omnibus (GEO; http://www.ncbi.nlm.nih.gov/geo/) under accession number GSE123981.

## SUPPLEMENTARY DATA

Anisimova_et_al_Supplemental_file.pdf.

Supplemental Methods. Sections 1–3. Detailed protocol of Ribo-Seq and RNA-Seq differential expression analyses. Section 4. Illustration of the analysis of age-related changes in transcript ribosomal coverage.

## Supplemental Figures (S1-S7)

Table S1. Ribo- and RNA-Seq library sizes and mapping percentage.

Table S2. Results of age-related gene expression and ribosome occupancy (RO) changes analysis.

Table S3. Ribo-Seq age-dependent GO enrichment analysis and ribosome occupancy (RO) GSEA.

Table S4. Putative transcriptional regulators of age co-expressed genes.

Table S5. Transcript isoform composition changes.

Table S6. Genes encoding mTOR-sensitive transcripts, 5’TOP containing transcripts and age-related changes in RO of genes encoding translation machinery.

## ETHICS APPROVAL

Experiments were carried out according to the protocols approved by the Institutional Animal Care and Use Committee (IACUC) of the Brigham and Women’s Hospital.

## ACKNOWLEDGEMENTS

We are grateful to Maria D. Logacheva, Genome Sequencing Research and Education Center, Faculty of Bioengineering and Bioinformatics MSU, for Illumina sequencing and valuable comments. We also thank Irina A. Eliseeva, Pavel V. Baranov, Dmitry E. Andreev and Nadezhda E. Makarova for discussion, and Ekaterina A. Sakharova, Alexander Tyshkovskiy and Philipp O. Gusev for assistance with bioinformatic analyses.

## ABBREVATIONS

CHX: cycloheximide
CPM: read counts per million
DE: differentially expressed
FDR: false discovery rate
GLM: generalized linear model
GO: Gene Ontology
GSEA: gene set enrichment analysis
PCA: principal component analysis
Ribo-Seq: ribosome profiling
RO: ribosome occupancy
TF: transcription factor
5’ TOP: 5’ terminal oligopyrimidine motif
TPM: transcripts per kilobase million
UTR: untranslated region.

## CONTRIBUTIONS

VNG, SED, ASA and MVG conceived the study; ASA and MVG carried out ribosome profiling and RNA-Seq; ASA, IVK and MBM performed computational analysis; VNG, SED, IVK and ASA interpreted the results; ASA, IVK, SED and VNG wrote the manuscript. All authors read and approved the final manuscript.

