## Supplementary data for "Multi-faceted deregulation of gene expression and protein synthesis with age"

**Content:**

Supplemental Methods:

Section 1. Ribo-Seq and RNA-Seq sequencing, data processing and bioinformatic analysis

Sections 2-4. Detailed protocol of Ribo-Seq and RNA-Seq differential expression analyses. Section 5. Illustration of the analysis of age-related changes in transcript ribosomal coverage.

Supplemental Figures:

Fig. S1 Analysis of RNA-Seq replicates of aging mouse liver and kidney.

Fig. S2 Representative examples of genes differentially expressed with age in mouse liver and kidney.

Fig. S3 Summary of genes differentially expressed with age identified in the study.

Fig. S4 Association of RO changes with changes in transcript isoform abundance.

Fig. S5 Decreased ribosome occupancy of transcripts encoding ribosomal and other translation-related proteins with age in mouse kidney.

Fig. S6 Metagene profiles of start and stop codon 5'ends according to ribosomal footprint coverage.

Fig. S7 Age-related gradual rearrangement of ribosome footprints towards the 3' end of coding sequence.

Supplemental references

### SUPPLEMENTAL METHODS

#### Section 1. Ribo-Seq and RNA-Seq sequencing, data processing and bioinformatic analysis

##### Ribo-Seq and RNA-Seq sequencing and data processing

Quality control of sequencing reads was performed with FastQC v0.11.5 (53). For Ribo-Seq data, the adapter sequences were cut with CutAdapt 1.14 (54), and quality trimming was performed with Sickle 1.33 (55). The Ribo-Seq and RNA-Seq reads were aligned to the mouse transcriptome and genome assemblies (mm10, downloaded from the UCSC Genome Browser using GENCODE M13 annotation. Alignment and basic read counting were performed with STAR 2.5.3 (56) with default parameters except for the number of allowed mismatches: up to 5% and 3% for Ribo-Seq and RNA-Seq, respectively, i.e. allowing approximately 1 mismatch per ~30nt (a typical Ribo-Seq footprint) and 2 per 75nt (an RNA-Seq read).

##### Ribo-Seq and RNA-Seq differential expression analysis

Raw gene counts processing and statistical analysis was performed in R Environment using edgeR Bioconductor package (57). RNA-Seq and Ribo-Seq data were RLE-normalized, separately for kidney and liver. Genes not reaching 1 read count per million (CPM) in at least one library were excluded from analysis resulting in 8,992 and 11,461 genes available for the analysis in liver and kidney, respectively.

To analyze age-dependent gene expression dynamics, we used Ribo-Seq data, as ribosome footprint counts provide combined data on both transcript abundance and translation efficiency. First, we used the generalized linear model (GLM, glmQLFit, and glmQLFTest of the edgeR package) with the age as a categorical variable (see Supplementary materials, Section 2). To identify differentially expressed genes, we compared gene expression in the studied ages with the expression data in organ samples of 3-month-old mice and considered genes to be differentially expressed if they pass the 0.05 Benjamini-Hochberg adjusted P-values threshold.  The number of DE genes was 689 in liver (excluding 1-month group) and 2,001 in kidney. To identify and characterize the groups of genes with similar patterns of age-dependent changes, the coexpression clusters, we clustered the genes using the expression fold change vectors (11, 20 and 26 versus 3 months) with ward.D clustering algorithm and Euclidean distance (Fig. 2A).  For up- and down-regulated genes we calculated Gene Ontology biological process (BP) and cellular compartment (CC) enrichment with the clusterProfiler package (58) (q-value threshold of 0.05, minimum gene set size of 2). The obtained GO enrichment data were visualized with REVIGO (59) (Fig. 2B). Second, we assessed a linear trend of age-dependent changes in gene expression (Fig. 3A). To this end, we used the edgeR GLM with an alternative design matrix where age as a predictor was considered a continuous variable (see Supplementary materials, Section 3). In all cases, genes were considered differentially expressed if the respective Benjamini-Hochberg adjusted P-values passed the 0.05 threshold.

**Principal component analysis of Ribo-Seq and RNA-Seq expression profiles**

To characterize RNA- and Ribo-Seq data, we applied principal component analysis (PCA) to 8,562 genes covered by more than one CPM in each sample of mouse liver and kidney Ribo- and RNA-Seq datasets (Fig. 1B). For this purpose, the gene counts were RLE normalized together for liver and kidney.

**Distribution of the Ribo-Seq and RNA-Seq coverage to the genome regions**

The Ribo-Seq and RNA-Seq genomic alignment coverage was mapped to the genome regions 5' and 3' untranslated regions (UTRs), coding sequences (CDS) and introns with read_distribution.py script from RSeQC-2.3.7 (60).

**Identifying putative transcriptional regulators of coexpressed genes**

For sets of up- and downregulated genes (Fig. 2A), we searched for potential transcriptional regulators and pairs of regulators with binding sites enriched at promoters and thus potentially involved in transcriptional response. To this end, we used the ChIP-Seq-based mouse cistrome data (61) for 315 transcription factors. As putative target genes of a particular transcription factor (TF), we considered those for which a cistrome segment with binding motif occurrences was within 500 nt from the transcription start site annotated in GENCODE. For each TF, we considered a 2x2 contingency table classifying genes by two binary features, being a putative TF target and being included in a particular gene list. Then, we applied right-sided Fisher's exact test and adjusted the resulting P-values using Benjamini-Hochberg correction for multiple tested TFs, separately for up- and down-regulated genes. We selected putative transcriptional regulators from the set of TFs passing adjusted P-values < 0.05 and odds ratios ~1.5 or more. The same analysis was performed for all pairs of TFs. To select the meaningful pairs, we used those of physically interacting TFs, based on the BioGRID dataset ALL-3.5.166 (62). Then, the sets of targets were intersected between the TFs forming the pair and used for the subsequent analysis.

**Analysis of transcript isoform composition changes**

To analyze changes in transcript isoform abundance, we compared shares of transcript isoform obtained with RSEM program (63) for young (3 months for kidney, 3 and 11 months for liver) and old animals (32 months for kidney, 26 and 32 months for liver) using Student's t-test with Benjamini-Hochberg correction for multiple tested genes. We found no particular genes passing 0.05 threshold for the adjusted P-value (Table S5). To visualize the relation between RO changes and changes in isoform abundance, we selected the genes with a mean share changing for more than 10% from total for at least one isoform and standard deviation within young and old groups being no more than 5% of the mean for kidney and 15% of the mean for liver. These genes were visualized among the list of genes sorted according to their linear RO Fold Change with age (Fig. S4, Tables S2 and S5).

**Ribosome occupancy analysis**

To assess the contribution of translation control in age-dependent gene expression changes, we estimated the transcript ribosome occupancy (RO, often referred to as the translation efficiency, TE). RO is often computed as the ratio of Ribo-Seq and RNA-Seq counts, which makes it non-trivial to estimate the statistical significance when comparing different time points. Here, we used edgeR GLM to estimate the contribution of Ribo-Seq relative to RNA-Seq with the following model formula: ~exp.type+age:exp.type, where exp.type is a categorical variable (Ribo-Seq or RNA-Seq) (see Supplementary materials, Section 4). The age was considered a continuous predictor variable. Such design not only allowed to subtract RNA-Seq contribution from Ribo-Seq read counts and thus obtain RO values, but also to detect age-dependent ribosome occupancy changes and assess their statistical significance by edgeR contrasts (61). Samples from 1-month-old mice were excluded from the analysis. Where indicated, paired comparison of RO between 32-months-old mice and 32-months old mice was used, with the age treated as categorical variable.

Functional enrichment of the groups of transcripts whose translation changed with age was determined for ranked gene lists sorted by their RO fold change with Gene Set Enrichment Analysis (GSEA) (64) implemented in fgsea R package (65) (Fig. 3B) and in Java desktop application (Fig. 4B). The GO terms with q-value less than 0.05 were then manually curated to form a non-redundant list.

**Construction of metagene profiles**

Metagene profiles of ribosomal footprints at the start and stop codons were built on Ribo-Seq transcriptome read alignment to reveal systematic age-induced changes in ribosomal coverage (Fig. 5A, Fig. 5SA). Transcript isoforms expression was estimated with RSEM (63). The major (the most expressed) isoforms of each gene reaching no less than 2.5 TPM (transcripts per kilobase million) in all libraries were selected for the subsequent analysis. The Ribo-Seq footprints coverage was estimated in 200nt windows centered at the start and stop codons using bedtools 2.27.0-1 (66). Liver (2,920 genes) and kidney (4,566 genes) data were analyzed separately, only transcripts with both start and stop codons annotated were selected. To construct the metagene profiles, for each gene, the coverage profile was normalized to the average Ribo-Seq transcript coverage and then summed up for all the genes.

**Analysis of ribosomal positional changes**

To analyze the dynamics of age-related changes in ribosomal coverage of protein-coding regions we, first, performed segmentation of transcript profiles to decrease the level of noise. We pooled ribosomal footprint coverage profiles for each age and replicate with an exception for 1 month-old mice samples and separately for organs and processed the pooled profiles with Poisson segmentation (Supplementary materials, Section 5) to split each transcript onto windows of different lengths, with stable ribosome footprints coverage within each window. Then, for each window, we calculated the log-ratio of the average ribosome footprint coverage at particular age to the mean coverage at 3 months using the list of transcripts selected previously for metagene profiles. This allowed to de-trend the profiles and construct a linear model for the ribosome footprint coverage with the relative transcript coordinate as a predictor variable (see Supplementary materials, Section 5). The distribution of linear regression slopes for each age was then visualized (Fig. 5C, Fig. S7) and the significance of shift between the distribution of slopes of different ages was assessed with the SIGN.test function from BSDA R package.

**Section 2. Ribo-Seq and RNA-Seq differential expression analysis.** Paired comparisons of Ribo-Seq gene expression in tissue samples of each age with samples of 3 months old mice. Age is taken as a categorical variable. R code extract (example for liver samples).

```
> groups
                             Name Organ Age Replicate Seqtype
ribo_liver_1m_1   ribo_liver_1m_1 liver   1         1    ribo
ribo_liver_1m_3   ribo_liver_1m_3 liver   1         3    ribo
ribo_liver_1m_2   ribo_liver_1m_2 liver   1         2    ribo
ribo_liver_3m_2   ribo_liver_3m_2 liver   3         2    ribo
ribo_liver_3m_1   ribo_liver_3m_1 liver   3         1    ribo
ribo_liver_3m_3   ribo_liver_3m_3 liver   3         3    ribo
ribo_liver_10m_1 ribo_liver_10m_1 liver  10         1    ribo
ribo_liver_10m_2 ribo_liver_10m_2 liver  10         2    ribo
ribo_liver_10m_3 ribo_liver_10m_3 liver  10         3    ribo
ribo_liver_20m_1 ribo_liver_20m_1 liver  20         1    ribo
ribo_liver_20m_2 ribo_liver_20m_2 liver  20         2    ribo
ribo_liver_20m_3 ribo_liver_20m_3 liver  20         3    ribo
ribo_liver_26m_1 ribo_liver_26m_1 liver  26         1    ribo
ribo_liver_26m_3 ribo_liver_26m_3 liver  26         3    ribo
ribo_liver_26m_2 ribo_liver_26m_2 liver  26         2    ribo
ribo_liver_32m_2 ribo_liver_32m_2 liver  32         2    ribo
ribo_liver_32m_3 ribo_liver_32m_3 liver  32         3    ribo

> design <- model.matrix(~Age, groups)
> design
                 (Intercept) Age1 Age10 Age20 Age26 Age32
ribo_liver_1m_1            1    1     0     0     0     0
ribo_liver_1m_3            1    1     0     0     0     0
ribo_liver_1m_2            1    1     0     0     0     0
ribo_liver_3m_2            1    0     0     0     0     0
ribo_liver_3m_1            1    0     0     0     0     0
ribo_liver_3m_3            1    0     0     0     0     0
ribo_liver_10m_1           1    0     1     0     0     0
ribo_liver_10m_2           1    0     1     0     0     0
ribo_liver_10m_3           1    0     1     0     0     0
ribo_liver_20m_1           1    0     0     1     0     0
ribo_liver_20m_2           1    0     0     1     0     0
ribo_liver_20m_3           1    0     0     1     0     0
ribo_liver_26m_1           1    0     0     0     1     0
ribo_liver_26m_3           1    0     0     0     1     0
```

```
ribo_liver_26m_2           1    0     0     0     1     0
ribo_liver_32m_2           1    0     0     0     0     1
ribo_liver_32m_3           1    0     0     0     0     1

> dgList_filtered_norm
An object of class "DGEList"

> dgList_filtered_norm <- estimateDisp(dgList_filtered_norm, design)
> fit <- glmQLFit(dgList_filtered_norm, design)

> qlf.1vs3 <- glmQLFTest(fit, coef=2)
> qlf.11vs3 <- glmQLFTest(fit, coef=3)
> qlf.20vs3 <- glmQLFTest(fit, coef=4)
> qlf.26vs3 <- glmQLFTest(fit, coef=5)
> qlf.32vs3 <- glmQLFTest(fit, coef=6)
```

**Section 2. Ribo-Seq and RNA-Seq differential expression analysis.** Analysis of linear age-dependent changes in gene expression according to Ribo-Seq data with the age taken as a continuous variable. R code extract (example for liver samples).

```
>   groups
                             Name Organ Age Replicate Seqtype
ribo_liver_3m_2   ribo_liver_3m_2 liver   3         2    ribo
ribo_liver_3m_1   ribo_liver_3m_1 liver   3         1    ribo
ribo_liver_3m_3   ribo_liver_3m_3 liver   3         3    ribo
ribo_liver_10m_1 ribo_liver_10m_1 liver  10         1    ribo
ribo_liver_10m_2 ribo_liver_10m_2 liver  10         2    ribo
ribo_liver_10m_3 ribo_liver_10m_3 liver  10         3    ribo
ribo_liver_20m_1 ribo_liver_20m_1 liver  20         1    ribo
ribo_liver_20m_2 ribo_liver_20m_2 liver  20         2    ribo
ribo_liver_20m_3 ribo_liver_20m_3 liver  20         3    ribo
ribo_liver_26m_1 ribo_liver_26m_1 liver  26         1    ribo
ribo_liver_26m_3 ribo_liver_26m_3 liver  26         3    ribo
ribo_liver_26m_2 ribo_liver_26m_2 liver  26         2    ribo
ribo_liver_32m_2 ribo_liver_32m_2 liver  32         2    ribo
ribo_liver_32m_3 ribo_liver_32m_3 liver  32         3    ribo

>   design <- model.matrix(~Age, groups)
>   design
                 (Intercept) Age
ribo_liver_3m_2            1   3
ribo_liver_3m_1            1   3
ribo_liver_3m_3            1   3
ribo_liver_10m_1           1  10
ribo_liver_10m_2           1  10
ribo_liver_10m_3           1  10
ribo_liver_20m_1           1  20
ribo_liver_20m_2           1  20
```

```
ribo_liver_20m_3           1  20
ribo_liver_26m_1           1  26
ribo_liver_26m_3           1  26
ribo_liver_26m_2           1  26
ribo_liver_32m_2           1  32
ribo_liver_32m_3           1  32
```

```
>   dgList_filtered_norm
An object of class "DGEList"
```

```
>   dgList_filtered_norm <- estimateDisp(dgList_filtered_norm, design)
>   fit <- glmQLFit(dgList_filtered_norm, design)
>   qlf.cont.age <- glmQLFTest(fit)
```

**Section 4. Analysis of mRNA ribosome occupancy (RO) linear changes with age.** Contribution of Ribo-Seq to gene expression is assessed relative to RNA-Seq. Age is taken as a continuous variable and the type of data (Ribo-Seq or RNA-Seq) is taken as a categorical variable. R code extract (example for liver samples).

```
> groups
                              Name Organ Age Replicate Seqtype
ribo_liver_3m_2   ribo_liver_3m_2 liver   3         2    ribo
ribo_liver_3m_1   ribo_liver_3m_1 liver   3         1    ribo
ribo_liver_3m_3   ribo_liver_3m_3 liver   3         3    ribo
ribo_liver_10m_1 ribo_liver_10m_1 liver  10         1    ribo
ribo_liver_10m_2 ribo_liver_10m_2 liver  10         2    ribo
ribo_liver_10m_3 ribo_liver_10m_3 liver  10         3    ribo
ribo_liver_20m_1 ribo_liver_20m_1 liver  20         1    ribo
ribo_liver_20m_2 ribo_liver_20m_2 liver  20         2    ribo
ribo_liver_20m_3 ribo_liver_20m_3 liver  20         3    ribo
ribo_liver_26m_1 ribo_liver_26m_1 liver  26         1    ribo
ribo_liver_26m_3 ribo_liver_26m_3 liver  26         3    ribo
ribo_liver_26m_2 ribo_liver_26m_2 liver  26         2    ribo
ribo_liver_32m_2 ribo_liver_32m_2 liver  32         2    ribo
ribo_liver_32m_3 ribo_liver_32m_3 liver  32         3    ribo
rna_liver_3m_2     rna_liver_3m_2 liver   3         2     rna
rna_liver_3m_1     rna_liver_3m_1 liver   3         1     rna
rna_liver_3m_3     rna_liver_3m_3 liver   3         3     rna
rna_liver_10m_1   rna_liver_10m_1 liver  10         1     rna
rna_liver_10m_2   rna_liver_10m_2 liver  10         2     rna
rna_liver_10m_3   rna_liver_10m_3 liver  10         3     rna
rna_liver_20m_1   rna_liver_20m_1 liver  20         1     rna
rna_liver_20m_2   rna_liver_20m_2 liver  20         2     rna
rna_liver_20m_3   rna_liver_20m_3 liver  20         3     rna
rna_liver_26m_1   rna_liver_26m_1 liver  26         1     rna
```

```
rna_liver_26m_3   rna_liver_26m_3 liver  26          3     rna
rna_liver_26m_2   rna_liver_26m_2 liver  26          2     rna
rna_liver_32m_2   rna_liver_32m_2 liver  32          2     rna
rna_liver_32m_3   rna_liver_32m_3 liver  32          3     rna

> design <- model.matrix(~Seqtype+Age:Seqtype, groups)
> design
                  (Intercept) Seqtyperibo Seqtyperna:Age Seqtyperibo:Age
ribo_liver_3m_2             1           1              0               3
ribo_liver_3m_1             1           1              0               3
ribo_liver_3m_3             1           1              0               3
ribo_liver_10m_1            1           1              0              10
ribo_liver_10m_2            1           1              0              10
ribo_liver_10m_3            1           1              0              10
ribo_liver_20m_1            1           1              0              20
ribo_liver_20m_2            1           1              0              20
ribo_liver_20m_3            1           1              0              20
ribo_liver_26m_1            1           1              0              26
ribo_liver_26m_3            1           1              0              26
ribo_liver_26m_2            1           1              0              26
ribo_liver_32m_2            1           1              0              32
ribo_liver_32m_3            1           1              0              32
rna_liver_3m_2              1           0              3               0
rna_liver_3m_1              1           0              3               0
rna_liver_3m_3              1           0              3               0
rna_liver_10m_1             1           0             10               0
rna_liver_10m_2             1           0             10               0
rna_liver_10m_3             1           0             10               0
rna_liver_20m_1             1           0             20               0
rna_liver_20m_2             1           0             20               0
rna_liver_20m_3             1           0             20               0
rna_liver_26m_1             1           0             26               0
rna_liver_26m_3             1           0             26               0
rna_liver_26m_2             1           0             26               0
rna_liver_32m_2             1           0             32               0
rna_liver_32m_3             1           0             32               0

> dgList_filtered_norm
An object of class "DGEList"

> dgList_filtered_norm <- estimateDisp(dgList_filtered_norm, design)
> fit <- glmQLFit(dgList_filtered_norm, design)
> qlf.cont.age <- glmQLFTest(fit, contrast=c(0,0,-1,1))
```

**Section 5.** Analysis of age-related changes in ribosomal footprint coverage of CDS (coding segments).

To analyze ribosome occupancy along the transcripts, we used the following workflow illustrated by ENSMUST00000070642.3 Cebpb transcript using the liver Ribo-Seq data (data for liver and kidney were considered separately).

**Panel A:** Ribo-Seq full-length footprint coverage normalized to total transcript coverage (3 months - orange, 32 months - blue, dashed lines represent the start and stop codons).

**A.**

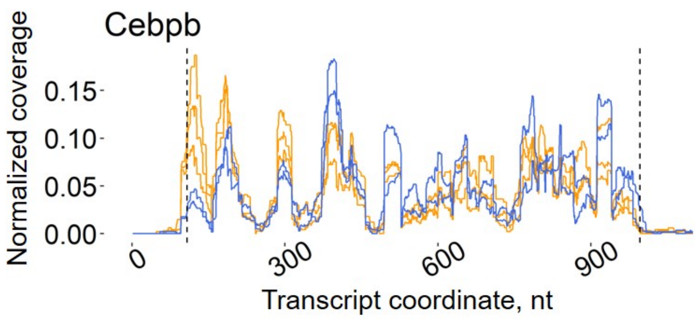


**Panel B:** Ribosome footprint coverage profiles were pooled across all age and replicates excluding 1-month-old mice samples.

**B.**

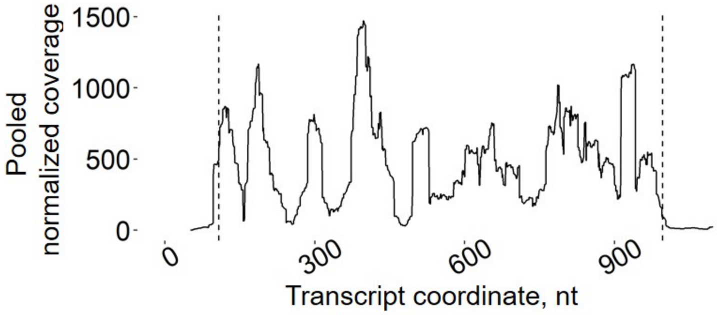


The pooled profiles were split into optimal segments of different lengths with *pasio* (https://github.com/autosome-ru/pasio, inspired by (67)) parameters alpha = 1 and beta = 1. This allowed obtaining a set of non-overlapping windows resembling the non-uniform distribution of ribosome footprints along each transcript.

For the further analyses we used the transcript CDS in two variants: either considering all segments including (-/+ 14nt) or by excluding (+/- 42nt) start and stop codons.

**Panel C:** The pooled profile processed with *pasio* with segments in the vicinity of the start and stop codons highlighted in green and red respectively.

**C.**

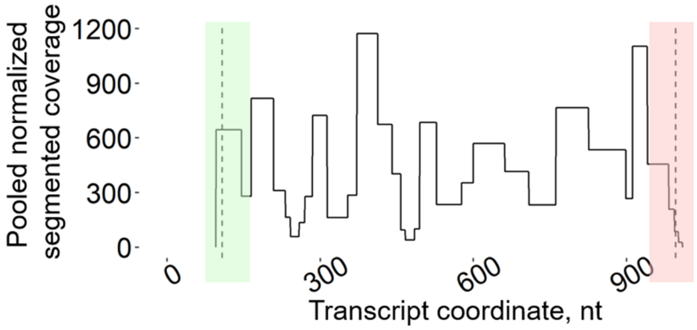


The pooled segmented profile provided a set of non-overlapping windows along the transcript characterized by particular ribosome occupancies. Next, for each library, we computed the average coverage of each segment and normalized the resulting profiles to the total coverage of a particular transcript (**Panel D**, 3- (orange) and 32-month-old (blue) mice samples). Pseudocount of 1 was added to the coverage value of each segment.

**D.**

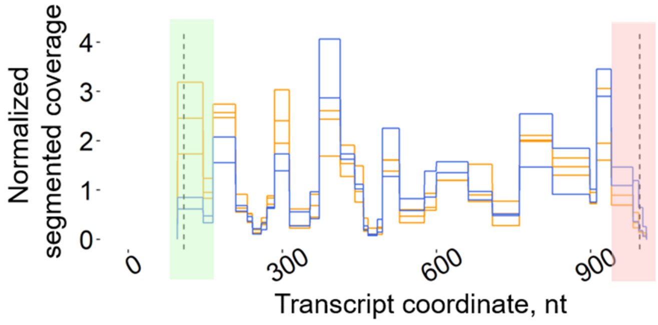


Next, we calculated the log ratio of segmented ribosomal footprint coverage at each age versus the mean coverage profile at 3 months (**Panel E**, 3- (orange) and 32-month-old (blue) mice samples). We considered all transcripts previously used for metagene profile construction.

The procedure allowed to obtain de-trended profiles (**Panel E**, each point shows an average coverage of a particular segment in a particular sample). The transcript coordinates (in nucleotides) of the segmented profile were projected into [0:1].

Next, for each transcript, we aggregated values across replicates and fitted a linear model of coverage depending on the relative transcript coordinate. The fitted linear models for 3- (orange) and 32-month-old (blue) mice are shown in the **panel E** as the solid lines. Distribution of regression slope for each age is shown in Fig. 5C and Fig. S7.

**E.**

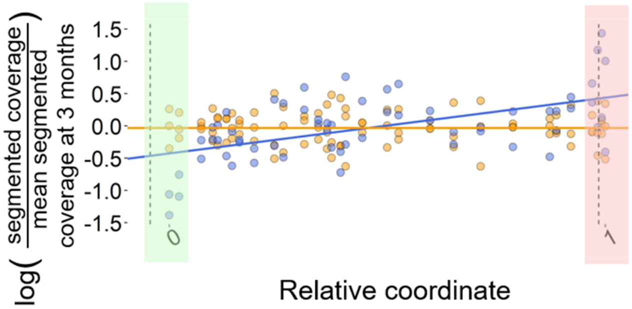


Results of linear regression analysis for ENSMUST00000070642.3 Cebpb transcript including start and stop codons are shown in **panel F**.

**F.**

| Age | Term | Slope | Std. error | Statistic | P-value |
|---|---|---|---|---|---|
| 3 | Intercept | -0.028 | 0.046 | -0.618 | 5.38E-01 |
| 3 | x | -0.004 | 0.080 | -0.054 | 9.57E-01 |
| 32 | Intercept | -0.427 | 0.096 | -4.426 | 4.12E-05 |
| 32 | x | 0.866 | 0.170 | 5.093 | 3.75E-06 |

SUPPLEMENTAL FIGURES

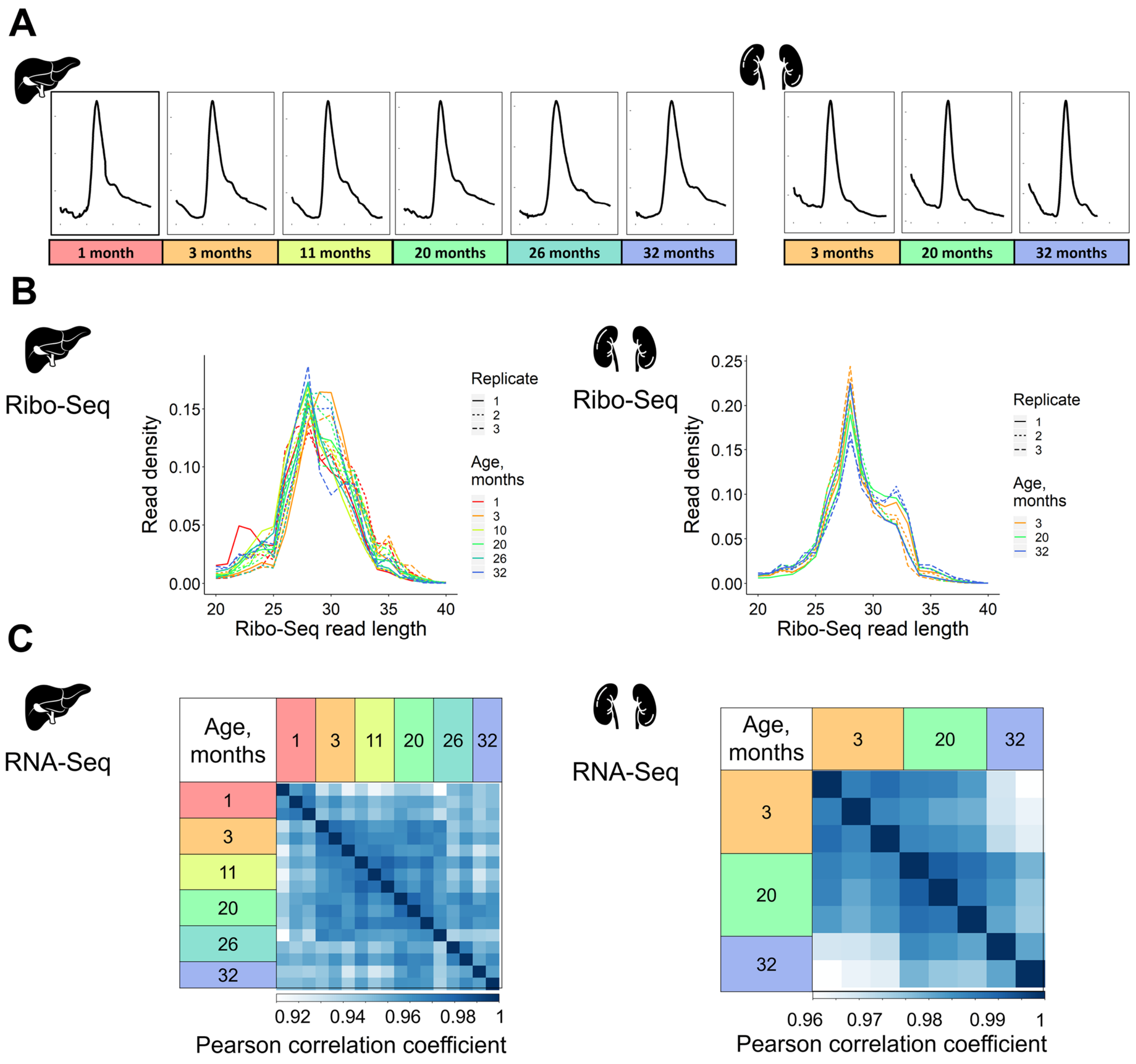


**Figure S1. Quality and reproducibility of Ribo-Seq and RNA-Seq.** (**A**) Monosome peaks after treatment of the samples with RNA digestion with RNase T1 and S7. (**B**) Ribo-Seq read length distribution. (**C**) Pearson correlation coefficient matrices of mouse liver (8,992 genes) and kidney (11,461 genes) RNA-Seq data.

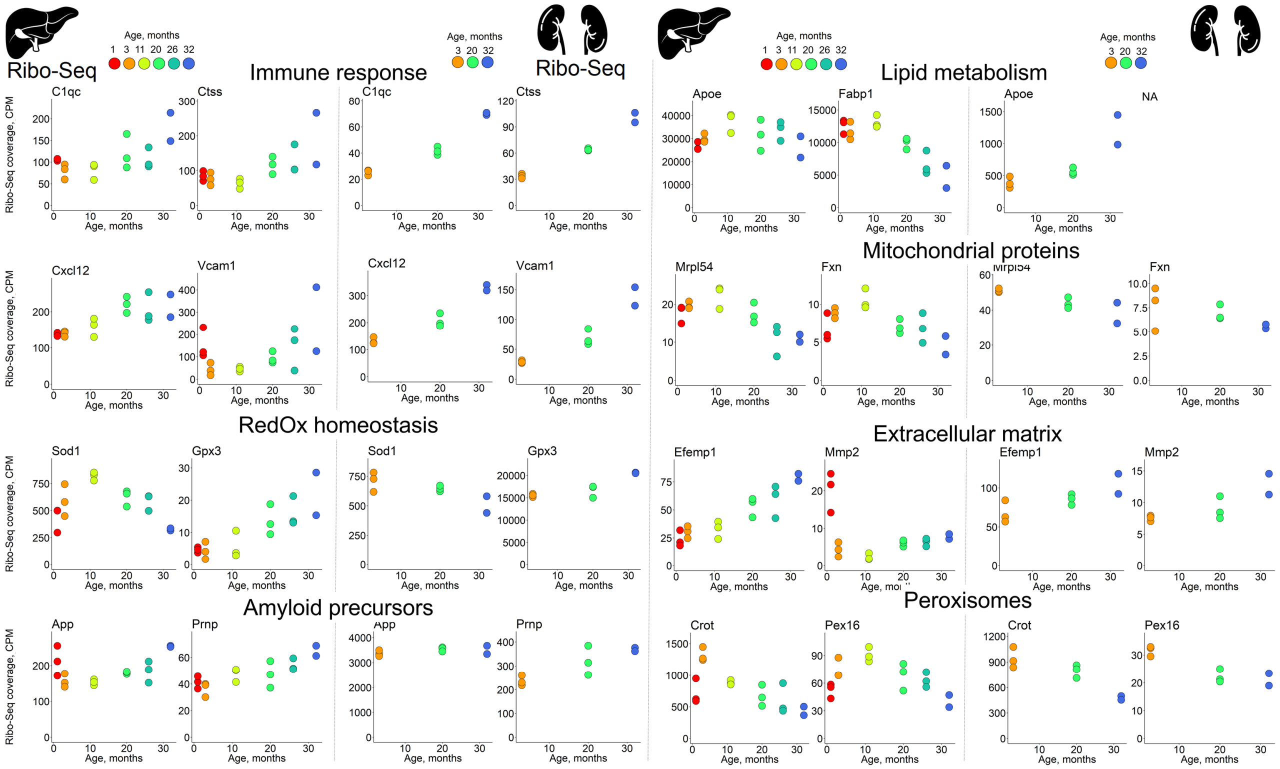


**Figure S2. Representative examples of genes differentially expressed with age in mouse liver and kidney.** Age-related patterns of ribosome profiling counts of genes belonging to functional groups found to be associated with aging in liver and kidney.

**A**

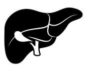

| From 8992 genes analyzed | | Differentially expressed for each age vs 3m | | | | | Linearly changed from 9110 genes analyzed |
|---|---|---|---|---|---|---|---|
| | Months / Direction | 1 | 11 | 20 | 26 | 32 | |
| Ribo-Seq | Up | 347 | 26 | 13 | 27 | 442 | 369 |
| | Down | 296 | 7 | 2 | 10 | 217 | 149 |
| RNA-Seq | Up | 259 | 56 | 21 | 28 | 126 | 211 |
| | Down | 295 | 15 | 4 | 9 | 48 | 80 |

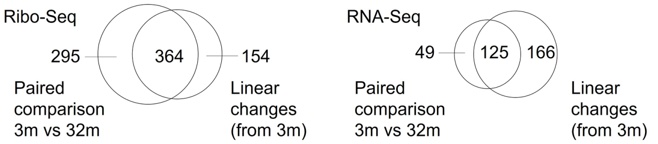


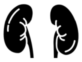

| From 11461 genes analyzed | | Differentially expressed for each age vs 3m | | Linearly changed from 11461 genes analyzed |
|---|---|---|---|---|
| | Months / Direction | 20 | 32 | |
| Ribo-Seq | Up | 199 | 1006 | 920 |
| | Down | 95 | 948 | 820 |
| RNA-Seq | Up | 134 | 477 | 610 |
| | Down | 90 | 394 | 556 |

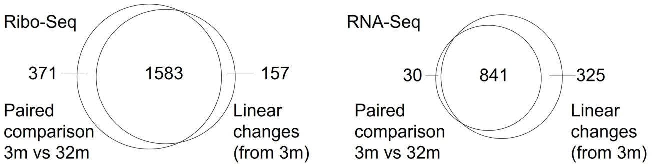


**B**

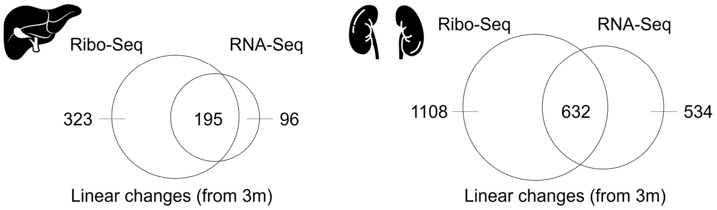


**Figure S3. Summary of genes differentially expressed with age identified in the study. (A)** Tables summarize the number of differentially expressed genes in Ribo-Seq and RNA-Seq data in paired comparisons of every age to 3 months and genes linearly changed from 3 to 32 months. **(B)** Venn diagrams illustrate the intersections between gene sets identified with linear and paired differential expression analysis.

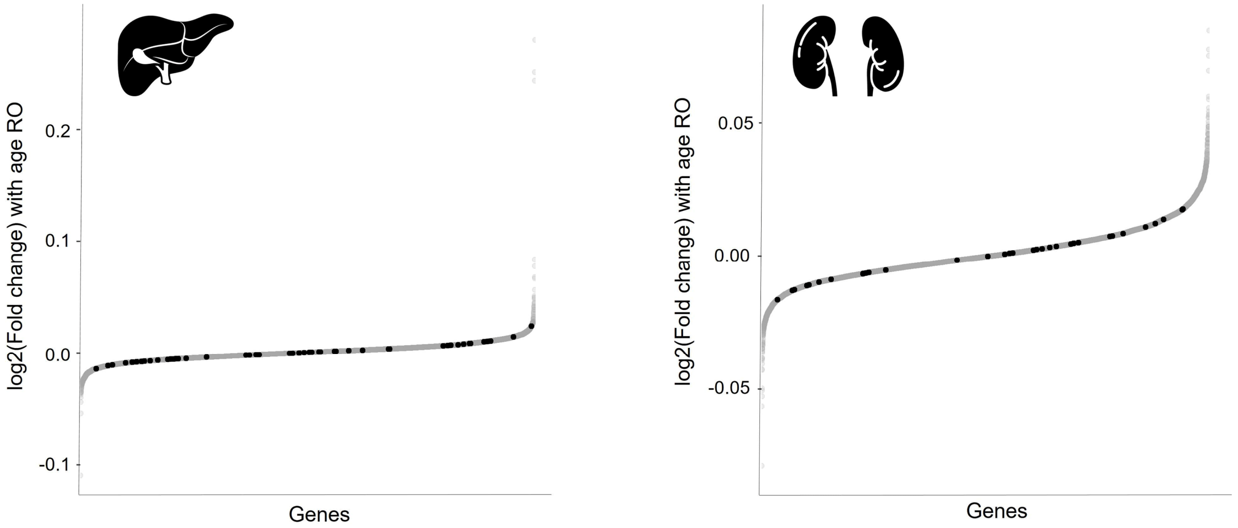


**Figure S4. Association of RO changes with changes in transcript isoform abundance.**

Genes with isoform abundance changed with age visualized (black) among the list of genes sorted according to their linear RO Fold Change with age (Tables S2 and S5). Genes were considered to exhibit age-related isoform abundance changes if the mean percentage of at least one of their isoforms changed more than 10% and standard deviation within young and old groups was no more than 5% of the mean for kidney and 15% for liver.

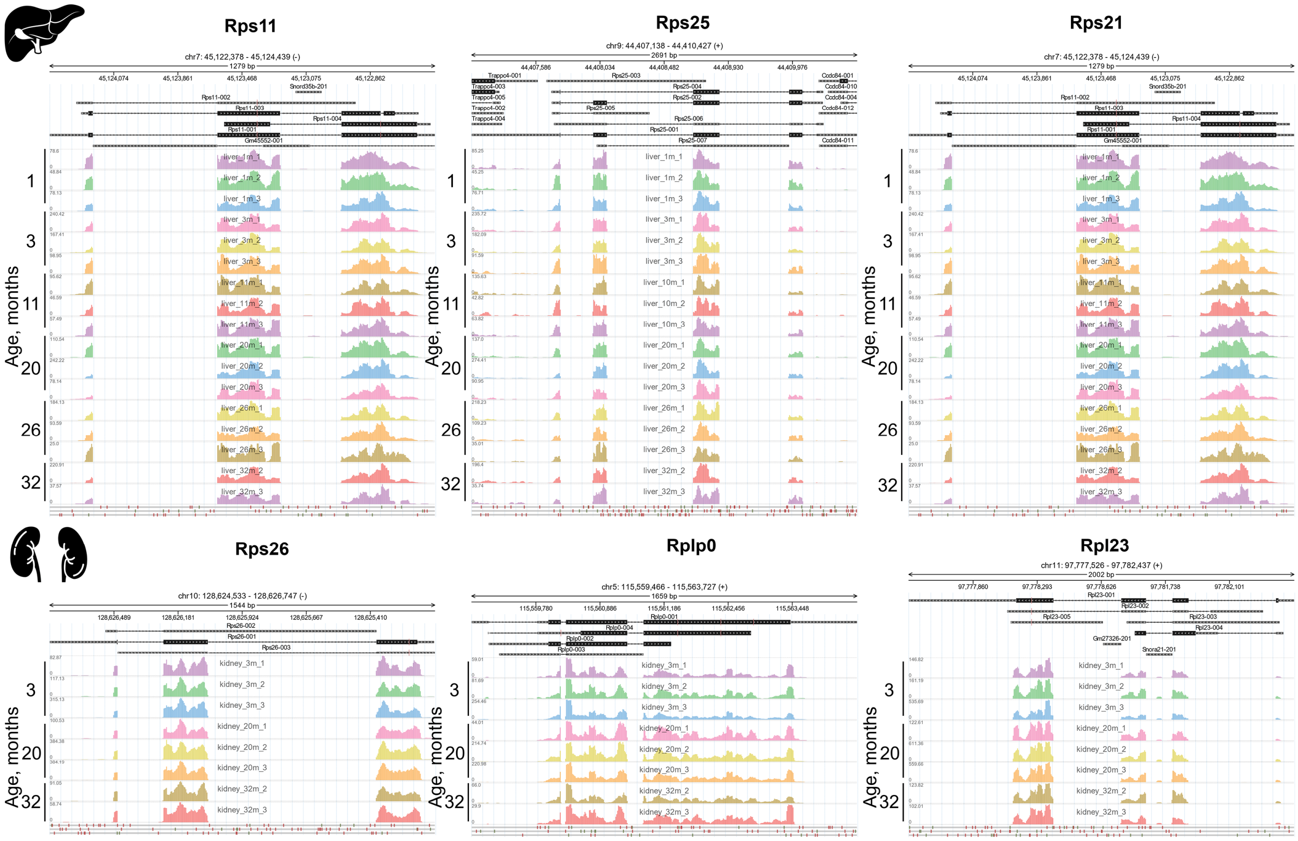


**Figure S5. Ribosomal protein encoding genes with the highest ribosome occupancy changes with age do not show isoform switch with age.** Ribo-Seq read coverage for the ribosomal protein encoding genes plotted with svist4get program (68).

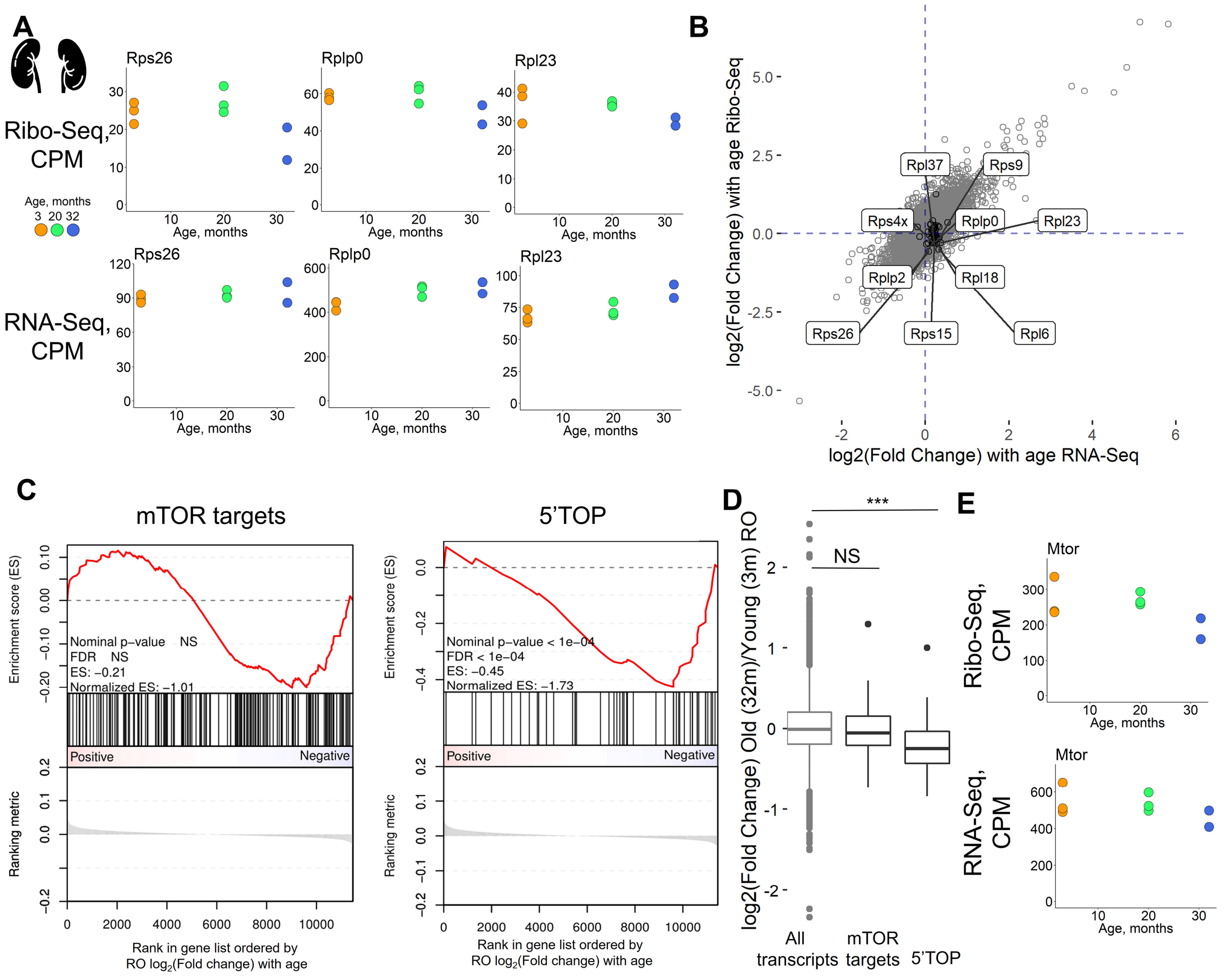


**Figure S6. Decreased ribosome occupancy of transcripts encoding ribosomal and other translation-related proteins with age in mouse kidney.** The decrease in ribosome occupancy of transcripts encoding ribosomal and other translation-related proteins with age. (**A**) Examples of age-dependent changes in Ribo- and RNA-Seq counts of genes coding for proteins with the functions associated with translation. (**B**) Comparison of transcriptome (RNA-Seq) and translation output (Ribo-Seq) of 32-months-old mice to 3-months-old-mice. Black dots represent 5'TOP genes. (**C**) GSEA of ribosome occupancy age-related changes (log2(Fold change), linear from 3- to 32 months old mice) in kidney of 44 5' TOP and 176 mTOR-sensitive genes (43). (**D**) Box plot showing distribution of mTOR regulated and 5'TOP genes ROs. Statistical significance was calculated with Mann-Whitney test. **E.** Ribo- and RNA-Seq counts of the *Mtor* gene.

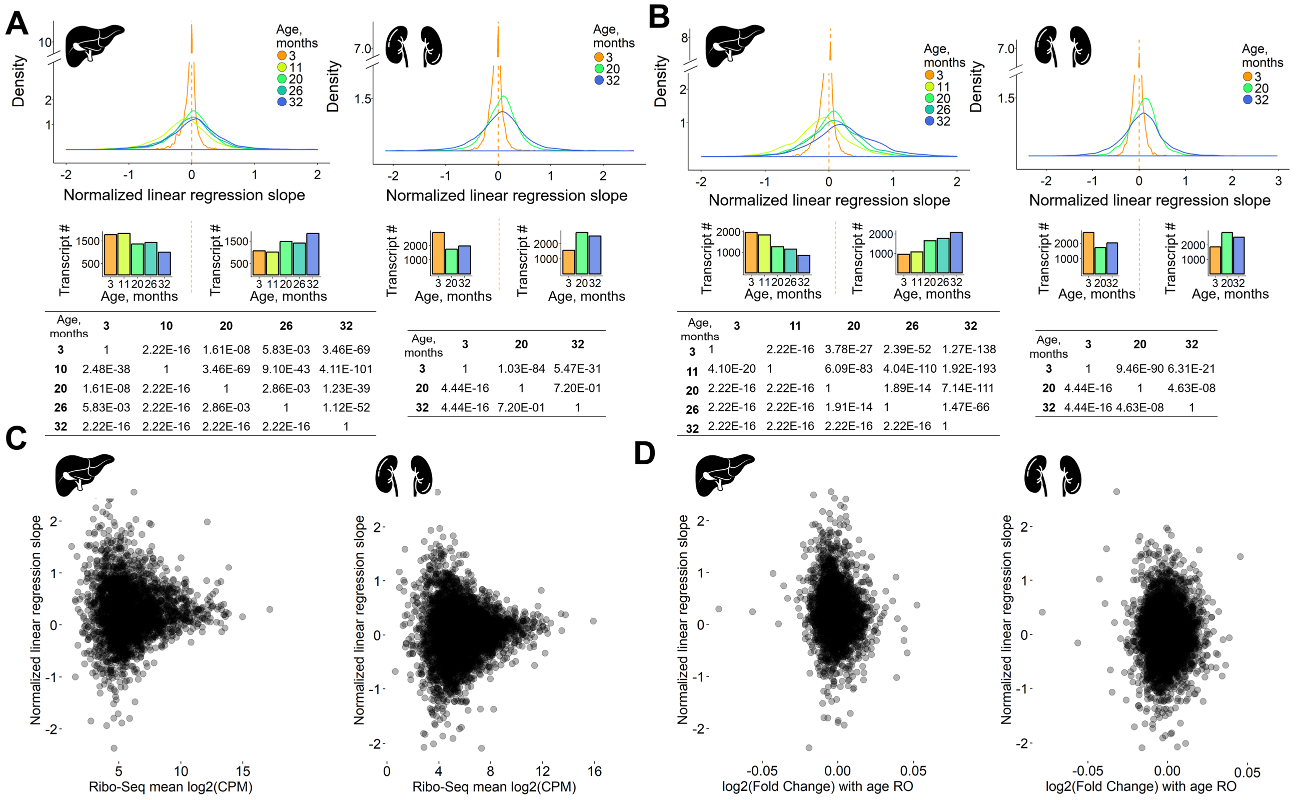


| Age, months | 3 | 10 | 20 | 26 | 32 |
|---|---|---|---|---|---|
| **3** | 1 | 2.22E-16 | 1.61E-08 | 5.83E-03 | 3.46E-69 |
| **10** | 2.48E-38 | 1 | 3.46E-69 | 9.10E-43 | 4.11E-101 |
| **20** | 1.61E-08 | 2.22E-16 | 1 | 2.86E-03 | 1.23E-39 |
| **26** | 5.83E-03 | 2.22E-16 | 2.86E-03 | 1 | 1.12E-52 |
| **32** | 2.22E-16 | 2.22E-16 | 2.22E-16 | 2.22E-16 | 1 |

| Age, months | 3 | 20 | 32 |
|---|---|---|---|
| **3** | 1 | 1.03E-84 | 5.47E-31 |
| **20** | 4.44E-16 | 1 | 7.20E-01 |
| **32** | 4.44E-16 | 7.20E-01 | 1 |

| Age, months | 3 | 11 | 20 | 26 | 32 |
|---|---|---|---|---|---|
| **3** | 1 | 2.22E-16 | 3.78E-27 | 2.39E-52 | 1.27E-138 |
| **11** | 4.10E-20 | 1 | 6.09E-83 | 4.04E-110 | 1.92E-193 |
| **20** | 2.22E-16 | 2.22E-16 | 1 | 1.89E-14 | 7.14E-111 |
| **26** | 2.22E-16 | 2.22E-16 | 1.91E-14 | 1 | 1.47E-66 |
| **32** | 2.22E-16 | 2.22E-16 | 2.22E-16 | 2.22E-16 | 1 |

| Age, months | 3 | 20 | 32 |
|---|---|---|---|
| **3** | 1 | 9.46E-90 | 6.31E-21 |
| **20** | 4.44E-16 | 1 | 4.63E-08 |
| **32** | 4.44E-16 | 4.63E-08 | 1 |

**Figure S7. Age-related gradual rearrangement of ribosome footprints towards the 3' end of coding sequence.** (**A**, **B**) Distribution of linear regression slopes for the normalized, segmented and smoothed transcript profiles pooled for each age depending on the relative transcript coordinate, excluding (**A**) or including (**B**) start and stop codons. The log ratio was taken between segmented ribosomal footprint coverage at each age and the mean coverage at 3 months for transcripts selected previously for metagene profile construction (see Supplementary Materials, Section 4). Bar plots depict the number of transcripts with negative (left) and positive (right) linear regression slope. Matrices summarize P-values of the sign test comparing differences of transcript linear regression slopes (slopes for the ages in rows were extracted from the slopes of the ages in columns). (**C**, **D**) Comparison between the extent of ribosome footprint rearrangement along the transcript with age (normalized linear regression slopes) with (**C**) mean gene Ribo-Seq coverage and (**D**) with ribosome occupancy (RO) linear changes with age (from 3 to 32 months).
